# Population-level integration of single-cell datasets enables multi-scale analysis across samples

**DOI:** 10.1101/2022.11.28.517803

**Authors:** Carlo De Donno, Soroor Hediyeh-Zadeh, Marco Wagenstetter, Amir Ali Moinfar, Luke Zappia, Mohammad Lotfollahi, Fabian J. Theis

**Affiliations:** Institute of Computational Biology, Helmholtz Center Munich, Munich, Germany; School of Life Sciences Weihenstephan, Technical University of Munich, Munich, Germany; Wellcome Sanger Institute, Wellcome Genome Campus, Cambridge, UK; Department of Mathematics, Technical University of Munich, Munich, Germany

## Abstract

The increasing generation of population-level single-cell atlases with hundreds or thousands of samples has the potential to link demographic and technical metadata with high-resolution cellular and tissue data in homeostasis and disease. Constructing such comprehensive references requires large-scale integration of heterogeneous cohorts with varying metadata capturing demographic and technical information. Here, we present *single-cell population level integration (scPoli)*, a semi-supervised conditional deep generative model for data integration, label transfer and query-to-reference mapping. Unlike other models, scPoli learns both sample and cell representations, is aware of cell-type annotations and can integrate and annotate newly generated query datasets while providing an uncertainty mechanism to identify unknown populations. We extensively evaluated the method and showed its advantages over existing approaches. We applied scPoli to two population-level atlases of lung and peripheral blood mononuclear cells (PBMCs), the latter consisting of roughly 8 million cells across 2,375 samples. We demonstrate that scPoli allows atlas-level integration and automatic reference mapping with label transfer. It can explain sample-level biological and technical variations such as disease, anatomical location and assay by means of its novel sample embeddings. We use these embeddings to explore sample-level metadata, enable automatic sample classification and guide a data integration workflow. scPoli also enables simultaneous sample-level and cell-level analysis of gene expression patterns, revealing genes associated with batch effects and the main axes of between-sample variation. We envision scPoli becoming an important tool for population-level single-cell data integration facilitating atlas use but also interpretation by means of multi-scale analyses.

## Introduction

The advancements in both experimental and computational single-cell technologies in recent years have enabled the generation of large-scale datasets comprising information from millions of cells. These datasets, also called “reference atlases”, collect data from different conditions and individuals and offer precious insight into cellular processes and states in different scenarios. Consortia such as the Human Cell Atlas (HCA) [1] and HuBMAP [2] aim to generate organ-level and body-level reference atlases which in turn allow researchers to study human organs from development to aging in both homeostatic healthy samples and disease. A key possibility opened by these atlases are *meta-analyses* relating cell types and states with different conditions such as disease state and individual demographics and lifestyle metadata such as age or smoking status [3, 4]. This in turn enables an understanding of which factors explain variation in gene expression at the cell level.

Performing large-scale meta-analysis on a single-cell atlas requires learning a joint representation of all datasets correcting for batch effects between them [5–7]. Tremendous efforts have been made in recent years to solve the data integration problem for single-cell RNA sequencing (scRNA-seq) datasets using a variety of modeling approaches ranging from statistical [8–11] and graph-based [12–14] methods to more complex deep learning models [5, 15–17]. Nonetheless, data integration for scRNA-seq remains a challenging problem [18], especially in the case of many datasets with a big variety of technical and biological metadata. Moreover, most current models do not provide insight on why data integration might have failed and do not make use of prior knowledge (e.g. cell type annotations).

With the increasing availability of large-scale atlases, leveraging these datasets has become a core focus in the field of single-cell genomics. Many analyses can be accelerated by mapping newly-generated data (referred to as *query* data) on top of an integrated atlas. Algorithms for efficient use of reference atlases are known as single-cell *reference mapping* methods [19–21], which build upon data integration algorithms to rapidly update an existing reference atlas by integrating a new query dataset. Transferring information from the reference atlas to the query enables efficient cell-type annotation of the query cells, automatic identification of novel (sub) populations [20, 22] and unseen disease or treatment affected cells [4, 22, 23].

Existing top performing deep learning reference building methods [6] rely on discrete, one-hot-encoded vectors to represent the condition to integrate such as sample or donor [15, 24]. These one-hot-encoded vectors are then concatenated to the input. This encoding does not allow for straight-forward downstream interpretation of the effect of each sample on the mapping, and results in a linear increase of the number of trainable parameters with the number of conditions to integrate. This can in turn make portability and maintenance of the model more challenging in large-scale collaborative efforts. Additionally, in the presence of many unique conditional categories (e.g. large cohorts), the number of conditional inputs can be closer or equal to the number of gene expression measurements (usually subset to a few thousand highly variable genes) leading to the model ignoring either input and producing inaccurate data representation and integration [25]. Among current reference building methods [10, 12, 24, 26] only scANVI and Seurat v3 offer built-in cell-type classification coupled with a reference mapping algorithm [19, 21]. Yet, while they can integrate annotated data to extend the reference, this requires retraining, which can be time consuming and resource demanding. Model retraining might also require additional access to reference data which is not always possible due to data sharing restrictions. This significantly limits both the extendability and usability of reference atlases for both atlas builders and downstream users.

Here, we introduce *single-cell Population Level Integration* (*scPoli*), a semi-supervised conditional generative model [5, 27] combined with advances in meta-learning [28] which is able to learn rep-resentations for both cells and samples (or other batch covariates), referred to in this work as conditional or sample embeddings. scPoli thus offers both a cell-level and a sample-level view of the dataset allowing users to perform multi-scale analysis. We call multi-scale analysis the simultaneous exploration of sample-level and cell-level representations obtained with our model.

scPoli uses prototype-based label transfer at the cell level and is augmented with an uncertainty estimation mechanism which can identify unknown cell-types in the query data. We demonstrate that scPoli is competitive in data integration and cell-type annotation compared to state-of-the-art methods in both tasks across six datasets, while also offering novel sample-level representations. We further showcase the features of our model features by integrating 166 samples across 11 datasets of lung studies, and performing query-to-reference mapping for two queries. We show potential use cases of conditional embeddings such as sample classification and data integration workflow guidance. Finally, we build a reference consisting of roughly 7.8 million PBMCs from 2,375 samples and 33 datasets, and explore the rich sample-level space obtained with scPoli finding genes associated with the main axes of sample variation. scPoli builds upon the transfer-learning framework scArches [21] and can be used to perform query-to-reference mapping.

## Results

### scPoli learns joint cell and sample representations

The variation of gene expression (*x_i_*) in a dataset can be ascribed to undesirable batch effects and actual biological signals. Similarly to other conditional probabilistic models for single-cell data [15, 29], scPoli aims to regress out batch effects in a non-linear fashion by means of a conditional variable (*s_i_*) representing batch (e.g. samples or datasets), while retaining biologically meaningful information. Moreover, scPoli posits that cell identities (*c_i_*) can be represented with learnable cell-type prototypes [30] modeled using latent cell representations (*z_i_*) (see **Fig. 1a**). scPoli, therefore, introduces two main modifications to the standard conditional variational autoencoder (CVAE) architecture which constitutes the backbone of many successful methods addressing problems such as data integration [5, 15, 24] and perturbation modeling [16, 29] in single-cell genomics. These modifications are (i) the replacement of one-hot-encoded vectors with continuous vectors of fixed dimensionality to represent the conditional term, and (ii) the usage of cell type prototypes to enable label transfer.

**Figure 1:**
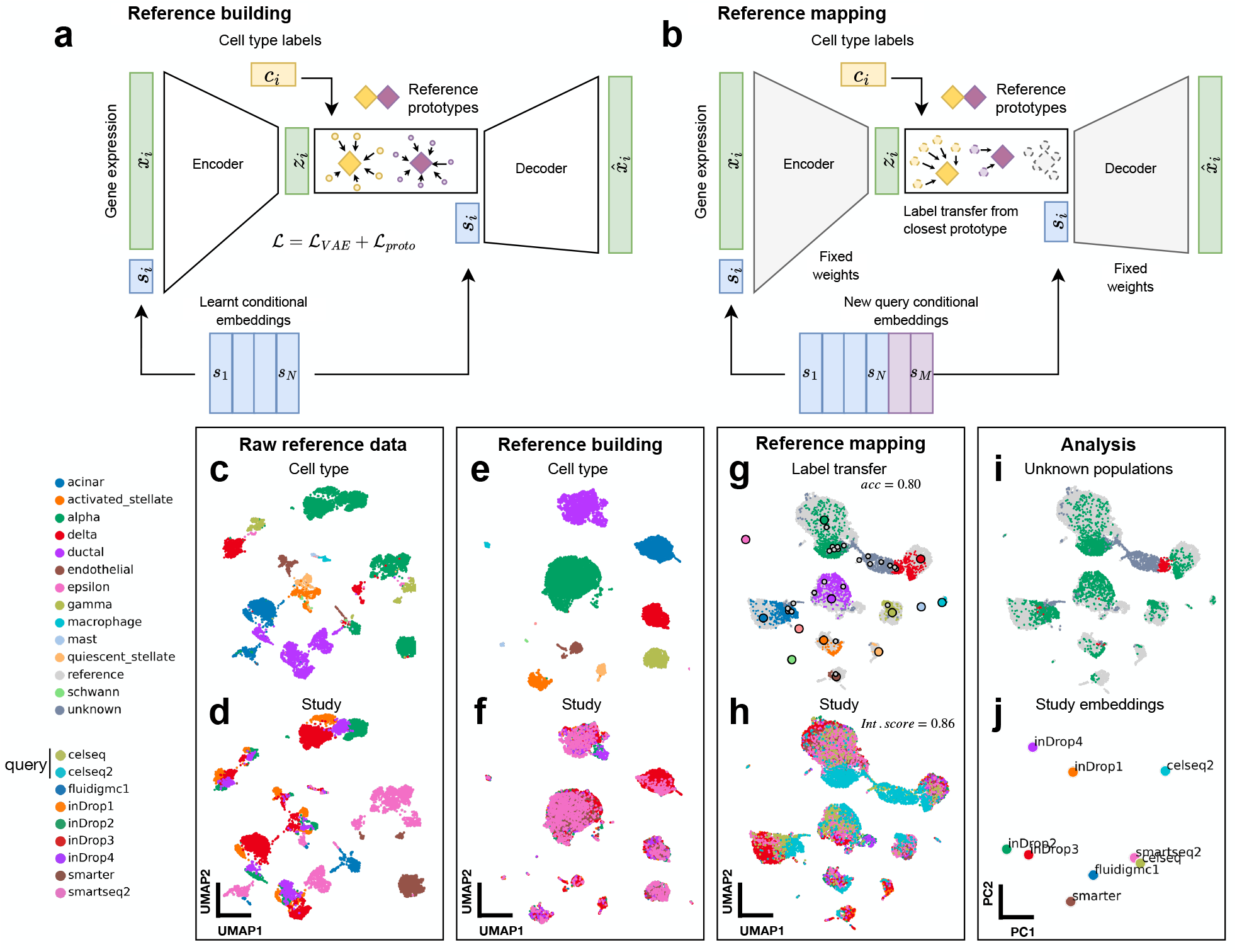
scPoli enables learning cell-level and sample-level representations. **(a)** scPoli reference building: the model integrates different datasets and learns conditional embeddings for each integrated study and a set of cell type prototypes. **(b)** scPoli reference mapping: the model weights are frozen (in grey) and a new set of conditional embeddings are added to the model. Cell type labels are transferred from the closest prototype in the latent space. Example of a standard workflow using scPoli on multiple pancreas datasets. **(c)** UMAP of the raw data to be integrated in a reference (13,093 cells), showing cell types and **(d)** studies by color. **(e, f)** Integrated reference data. **(g)**3,289 query cells (celseq and celseq2 studies) are projected onto the reference data in the reference mapping step. UMAPs show in color the query cells and in grey the reference cells. Reference cell type prototypes are shown in bigger circles with a black edge. The accuracy of the label transfer is 80%. **(h)** Cells are colored by study or origin after reference mapping. The model achieves a mean integration score of 0.86. **(i)** Outcome of the label transfer step from reference to query. **(j)** Principal component analysis (PCA) of the conditional embeddings learned by scPoli.

CVAE-based methods encode each condition by means of one-hot-encoded representations 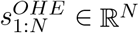, where *N* is the number of conditions. These are concatenated to the input, and an additional neuron for each condition is added to the first layer of the encoder neural network. The drawbacks of this approach become evident once N increases, which is becoming more common for single-cell atlases. In fact, in the case of thousands of conditions to be integrated, this approach leads to a notable increase in the number of total trainable parameters in the neural network architecture, which can slow down training. scPoli replaces these one-hot-encoded vectors with learnable conditional embeddings 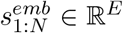 of fixed dimensionality that are associated with each conditional term (e.g. sample). These are concatenated to the input and learned at training time (**Fig. 1a**). Since the dimensionality of these embeddings does not depend on N, this approach offers more scalability in scenarios where many conditions are to be integrated. The number of trainable parameters is not a function of N anymore, and training speeds will be unaffected. Furthermore, unlike one-hot-encoded vectors, these learnable conditional embeddings capture meaningful representations of eacg integrated condition and can be visualized and analyzed, providing interesting insight in large-scale studies.

We designed scPoli as a reference building method integrated within our reference mapping technique scArches. scArches enables transfer learning from a reference model to analyze a query for conditional generative models employing architectural surgery[21]. After training a model on the reference dataset, the learned weights are frozen. Then, the query datasets are mapped by expanding the first layer of the encoder to accommodate the new *M* query conditions and adding a set of trainable weights from these new neurons towards the first hidden layer. The current implementation of architecture surgery requires deploying reference models together with an one-hot-encoder which needs to be adjusted every time a new query dataset is added. This can be problematic and reduces portability in the presence of many samples. In contrast, the architecture surgery for scPoli is done simply by freezing the weights of the model trained on the reference and learning a new set of M embeddings to accommodate the query data conditions (**Fig. 1b**).

The second novel addition to existing CVAE models is the incorporation of prototypes commonly used in meta-learning [31]. These allow efficient learning across tasks and datasets and have previously been used for cell-type classification in single-cell data [26]. scPoli models prototypes using the average gene expression of cells belonging to a cell-type and leverages them not only to transfer cell-type annotation in a partially labeled reference or in new query data, but also to improve data integration by introducing an additional term in the learning objective which we call prototype loss. This new term encourages the model to consolidate the latent representation of cells around their prototype, which we show leads to better preservation of biological signals after data integration (see **Methods**). Unlabeled cells are classified by comparing distances to the prototypes. The label of the closest prototype is then assigned as a predicted cell type label. We also exploit the distance of each cell to its closest prototype as a proxy for uncertainty in cell-type classification and identification of unknown cell-type and states available in the query data. Finally, prototypes enable extending an initial reference atlas with novel cell-types from a labeled query without retraining the reference model as opposed to existing methods [24].

We illustrate a standard scPoli workflow on a collection of 9 pancreas studies which have often been used to demonstrate data integration in other publications (see **Benchmark datasets**) in **Fig. 1c-j**. We build an integrated reference on 7 fully labeled datasets generated with multiple scRNA-seq protocols. After this step, we can assess that batch effects are removed while cell identities are preserved. We use two datasets (celseq, celseq2) as an unlabeled query and map them onto the reference data. After inspecting a joint UMAP of the query and reference datasets we can observe that query cells are mapped onto the reference data (mean integration score of 0.86) and that most cells are classified correctly with an accuracy of 80%. A cluster of cells (beta cells) that was removed from the reference dataset to mimic an unknown cell-type scenario is correctly identified. The conditional embeddings learned by scPoli can be visualized using any dimensionality reduction technique and capture similarities between the integrated samples. We use principal component analysis to keep linear relationships between the embeddings, and we can observe that all inDrop samples are mapped towards the same region in this embedding space.

### scPoli accurately integrates datasets and at the same time transfers cell-type annotations

To understand how well scPoli integrates different single-cell datasets, and how accurately it predicts cell type annotations, we benchmarked our model against a set of state-of-the-art methods for data integration and label transfer. Achieving a broad overview, we included in this comparison both deep learning models (scVI [32], scANVI [33] and MARS [26]) and other type of methods (Seurat v3 [34], Symphony [35] and a linear SVM). Out of these models, only our method, scANVI and Seurat v3 tackle both data integration and label transfer, while some exclusively do data integration (scVI, Symphony), and others only cell type classification (MARS, SVM). Moreover only scPoli, scVI and scANVI (the latter two via scArches) are capable of query-to-reference mapping, while the rest of the methods require the reference data and retraining in order to map a query.

All of these models, except for MARS and Symphony were part of Luecken *et al*. [6] data integration benchmark, where they came out as the consistent top performers, guiding our choice to pick them in this benchmark study.

We tested these methods on six datasets, spanning a variety of scenarios and technical properties (see **Benchmark datasets**) (**Supp. Fig.1)**. For each dataset a set of studies to use as reference was picked, while the rest was used as query. To quantify the performance of data integration we used metrics from the Luecken *et al*. benchmarking effort [6] (see **Methods**). We aimed to quantify how well the different methods preserved meaningful biological signal and how effective they were at correcting batch effects, therefore we included metrics for both aspects of the data integration problem. As done in the original benchmarking study, we obtained the overall score as a weighted average of the mean estimations of biological variation conservation and batch correction, with weights of 0.6 and 0.4 respectively.

We found that scPoli outperformed the next best performing model (scANVI) by 5.3% in data integration (**Fig. 2a**). When we looked separately at batch correction and biological conservation metrics, we observed that scPoli preserved biologically meaningful signals better than other methods (8.87% over scANVI), while achieving similar levels of batch correction. In order to understand whether the improvements stemmed from the use of conditional embeddings or by the inclusion of the prototype loss, we benchmarked two variants of our model. We included a scPoli model with standard one-hot-encoded vectors to represent batch, and a scPoli model trained without prototype loss. We found that the inclusion of the prototype loss to be the main driver of the improvement in biological conservation, with the scPoli model with one-hot-encodings achieving similar integration scores as scPoli. The scPoli model trained without prototype loss on the other hand showed improved batch correction scores at the sacrifice of biological conservation (**Fig. 2b**).

**Figure 2:**
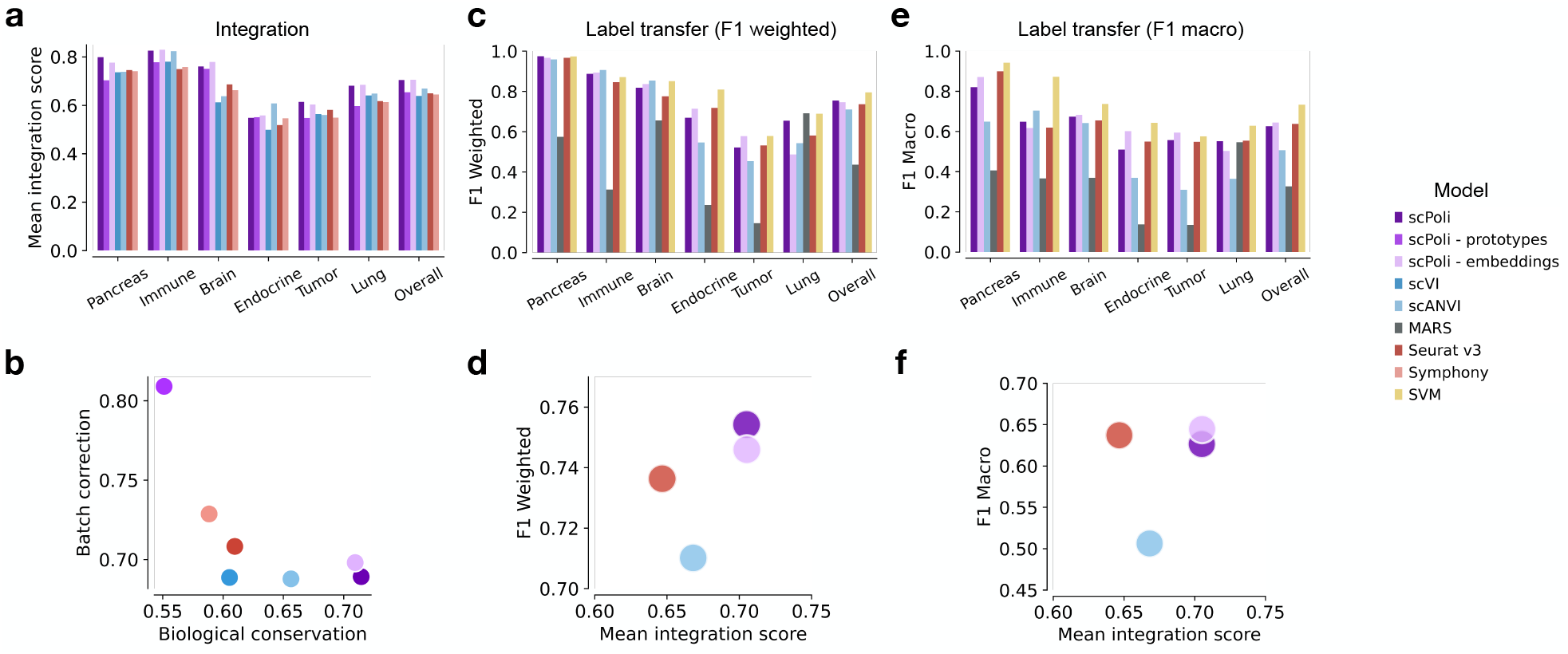
scPoli reaches state-of-the-art performance on data integration and label transfer. **(a)** Mean integration score obtained using the benchmarked models on different datasets. The barplots on the right show the average results across datasets. **(b)** Overall scores across datasets for biological conservation and batch correction performance of the benchmarked models. **(c)** Weighted F1 scores achieved by each model when classifying query cells on the various datasets. **(d)** Weighted F1 score and the overall integration score of the models capable of both data integration and label transfer. **(e)** Macro averaged F1 query classification scores achieved by each model on the various datasets. **(f)** Macro averaged F1 score and the overall integration score of the models capable of both data integration and label transfer.

To assess the quality of the classification achieved during the reference mapping step we used two metrics: the weighted average F1 score, and the macro averaged F1 score. When computing the weighted average F1 score, the score is first obtained for each cell type separately and then averaged weighted by the support of each class. This metric thus reflects how well each model performs on the most represented classes in the data. The macro averaged F1 score on the other hand, is computed by averaging each class’ F1 score without any weighting, therefore being more sensitive to the classification performance for the less represented cell types. We observed that scPoli outperformed all methods except for the linear SVM on the weighted F1 metric (**Fig. 2c**). Out of the models that are capable of both data integration and cell type classification scPoli came out on top (**Fig. 2d**). When looking at the macro averaged F1 score scPoli showed comparable performance to Seurat v3 and a sizeable improvement over scANVI, indicating better performance on less represented cell types (**Fig. 2e, f**).

### scPoli enables intepretable integration of the human lung cell atlas

We showcase the data integration capability and quality of label transfer yielded by scPoli by applying it on the Human Lung Cell Atlas (HLCA) [4]. The HLCA is a curated collection of 46 single-cell datasets of the human lung, with samples from 444 individuals. The atlas is divided into a core collection of data, which comprises data from 166 samples and 11 datasets, and an extended one which includes the remaining data including healthy and disease samples. The extended HLCA was annotated by the original authors by mapping samples onto the core reference and transferring labels. Following the work in the original study we used the HLCA core data for reference building. During this step we decided to integrate data at the sample level, rather than at the dataset level like in the original study. This choice was made to obtain a better resolution of the conditiona; embeddings and thus allow interpretation using more granular sample-level metadata. For prototype initialization and learning we used the finest level of annotations available, resulting in a set of 58 cell type prototypes.

When we visually inspected the UMAP of the integrated object, we observed that scPoli was able to successfully integrate the data from different studies (**Fig. 3 a**), while maintaining clear structure among the known cell identities (**Fig. 3 b**). We further assessed the quality of data integration and compared it against scANVI [24], which was selected as the best performing method in the HLCA integration task in the original publication. To keep the comparison consistent we also trained scANVI at the sample level. scPoli yielded an integration which preserved biologically meaningful variation better than scANVI, while achieving similar performance in batch correction (**Fig. 3 c**). This resulted in scPoli having an edge over scANVI in overall data integration performance.

**Figure 3:**
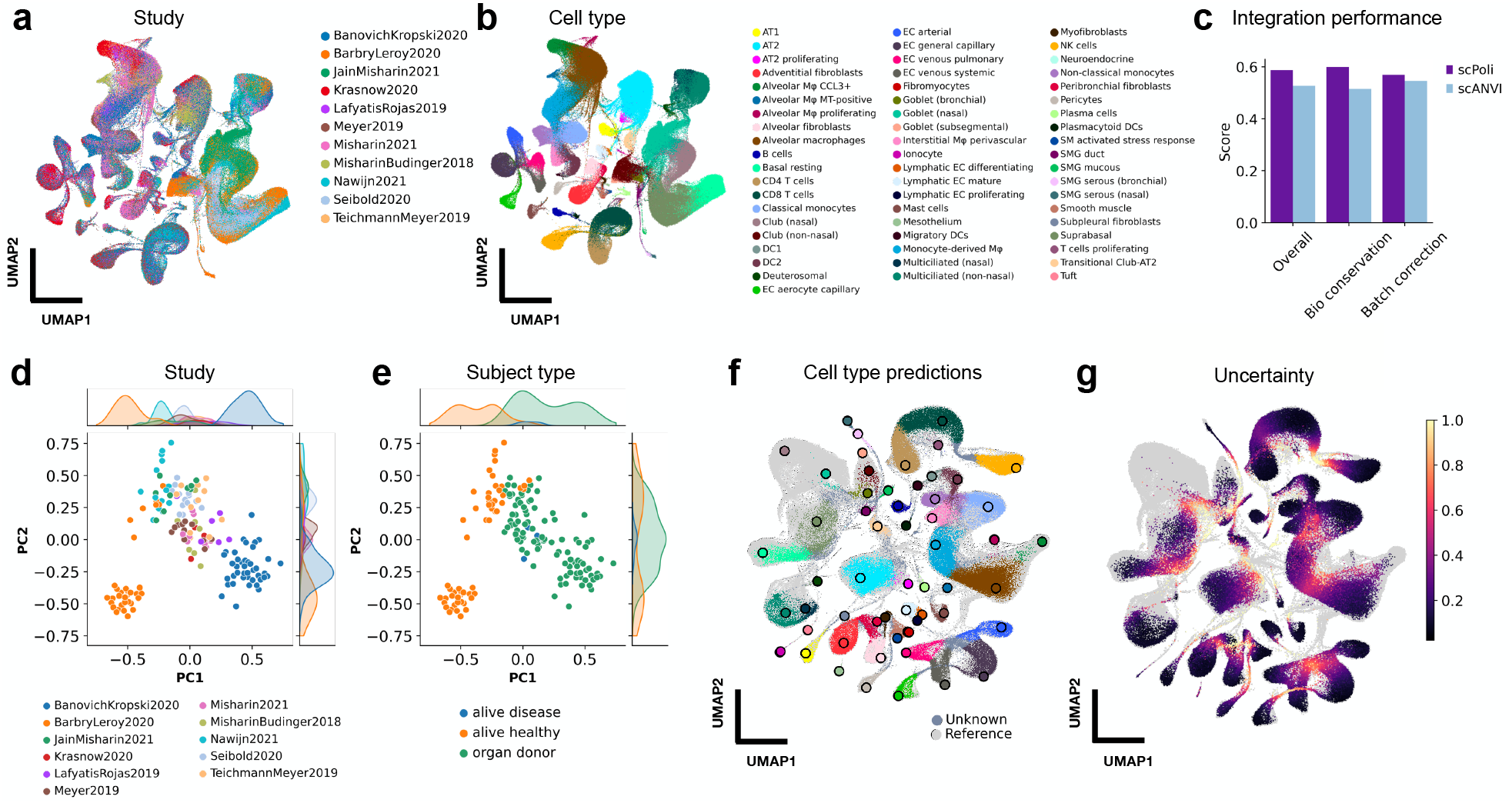
scPoli performs intepretable integration and query-to-reference mapping on the Human Lung Cell Atlas (HLCA) **(a)** UMAP of the integrated HLCA core after reference building, cells are color coded by their study of origin and **(b)** by cell type. **(c)** Comparison of integration performance yielded by scPoli and the scANVI. **(d)** Visualization of the first two PCs obtained with a principal component analysis of the sample embeddings learned from the reference data. Samples are color coded by their original study and **(e)** by sample type. **(f)** UMAP of the joint query and reference datasets after query-to-reference mapping for a healthy query. Reference cells are shown in light grey, query cells are colored by the predicted cell type and unknown cells are shown in dark grey. Legend for the predicted cells is shared with (b). Reference prototypes are shown as bigger dots with a black border, and are colored by cell type. **(f)** UMAP of the integrated object with uncertainties in color. Reference cells are shown in gray.

The use of learnable conditional embeddings instead of one-hot-encoded vectors allows our model to learn representations for each integrated condition that can be then visualized and analyzed. When looking at the first two principal components of the sample embeddings we found that samples from the same studies grouped together, reflecting likely similarities between data from the same dataset (**Fig. 3 d, Supp. Fig. 2 a**). The HLCA collects additional metadata associated with each sample including a variety of demographic and technical information. We analyzed how these covariates varied along this novel latent representation learned by our model. Interestingly, we found a set of covariates that showed meaningful variation across this latent representation. These included information regarding properties of the sample such as subject type (e.g. donor or alive) (**Fig. 3d**) and anatomical location (**Supp. Fig. 2 b**). Subject type refers to status of the subject from which the sample was taken: “alive” for biopsies from healthy or diseases subjects and “donor” for data collected from organs of diseased donors. Anatomical location refers to the anatomical origin of the collected tissue in the sample. Other covariates (e.g. sex or ethnicity) appeared to be mixed in the sample embedding space (**Supp. Fig. 2 c, d**), indicating that the main drivers behind batch effects between samples are likely to be related to the nature of the tissue, the way it was processed and other technical factors.

### scPoli enables query-to-reference mapping and label transfer

After building a reference model using the HLCA core dataset, we mapped a group of query samples of healthy patients from the HLCA extended dataset (Meyer *et al*., 2021). These data consist of six samples and contain 9 cell identities that are not present in the reference. In **Fig. 3 f** we show the result of the reference mapping and label transfer. As a proxy for uncertainty in cell type prediction we use the euclidean distance from the closest prototype in the reference, with the intuition that cells that fall far from any of these prototypes could represent an unknown or novel cell state. Similarly to the original HLCA study, in which a kNN-graph based uncertainty was used, we noticed that cells that lay in regions of transitions between cell types displayed the highest levels of uncertainty, as well as cells whose identities were not present in the core dataset used for reference building (**Fig. 3 g**). We considered all cells with an uncertainty higher than the 90% quantile as unknown (**Supp. Fig. 2 e**) and inspected the classification performance by cell type (**Supp. Fig. 2 g**). A subset of novel cells were successfully detected as unknown, especially chondrocytes, erythrocites and myelinating and non-myelinating Schwann cells. Natural killer T cells, Gamma-Delta T cells and regulatory T cells on the other hand were not detected as unknown, and were mostly classified as either CD4 T cells or CD8 T cells, which could also be a result of over-clustering in the original atlas [36]. Overall, scPoli achieved an accuracy of 75%, outperforming the model used in the original study, which yielded an accuracy of 69%. It is important to stress the fact that label transfer in scPoli happened without need for the reference dataset, resulting in a much more convenient and lean computation. The methodology used in the original HLCA paper requires the computation of a kNN graph on an integrated embedding of reference and query data which can be time-consuming and cumbersome when there are many cells. In contrast, scPoli transfers labels by comparing distances to a small set of prototypes that are obtained during the reference building step and stored within the reference model. This constitutes a big advantage in cases where the reference data cannot be shared due to privacy or computational reasons. Furthermore, we observed that scPoli is more robust at detecting unknown cells than the methodology involving a kNN graph and scANVI. We compared the ratio of true predictions (correct cell type prediction and correct unknown detection) across different thresholds for unknown cell type detection for three models and scPoli consistently obtained better accuracy compared to the other two methods (**Supp. Fig. 2 f**).

In order to see how scPoli would perform when mapping a query dataset from a different condition than the one in reference, we mapped data collected from cancer patients. These data contain two cell identities not present in the reference (cancer and erythrocytes) and should intuitively be harder to integrate due to the different condition introducing stronger batch effects. Nonetheless, we observed that scPoli mapped the query dataset successfully onto the reference (**Supp. Fig. 3 a**). Since this query has a much coarser cell type annotation, we mapped the labels obtained with scPoli to the cell types present in the query via an expert-curated mapping obtained from the authors of the original study. By inspecting the uncertainty distribution on a joint UMAP representation of the reference and query datasets, we observed that almost all cancer cells mapped to a large cluster of cells whose label prediction had high uncertainty and were classified as unknown by our model (**Supp. Fig. 3 b, c**). We broke down the classification accuracy of the model by cell type in the query dataset and observed that 85% of cancer cells and 98% of erythrocytes were identified as unknown as desired (**Supp. Fig. 3 d**).

### scPoli enables multi-scale classification of cells and samples

We tested unlabeled sample classification as a potential use-case for the novel conditional embeddings obtained with scPoli. We integrated a COVID-19 PBMC dataset by Su *et al*. [37] with large biological signals using the sample covariate as batch condition. The data contains 559,517 cells from 270 samples from 145 patients, across various states of COVID-19. We selected 30 random samples and set their cell type annotations and sample annotations as unknown. We then integrated the data and propagated the labels from the rest of the labeled dataset. We first assessed the quality of integration and label transfer which achieved an overall 90% accuracy on a coarse cell type annotation level (**Fig. 4 a, b**).

**Figure 4:**
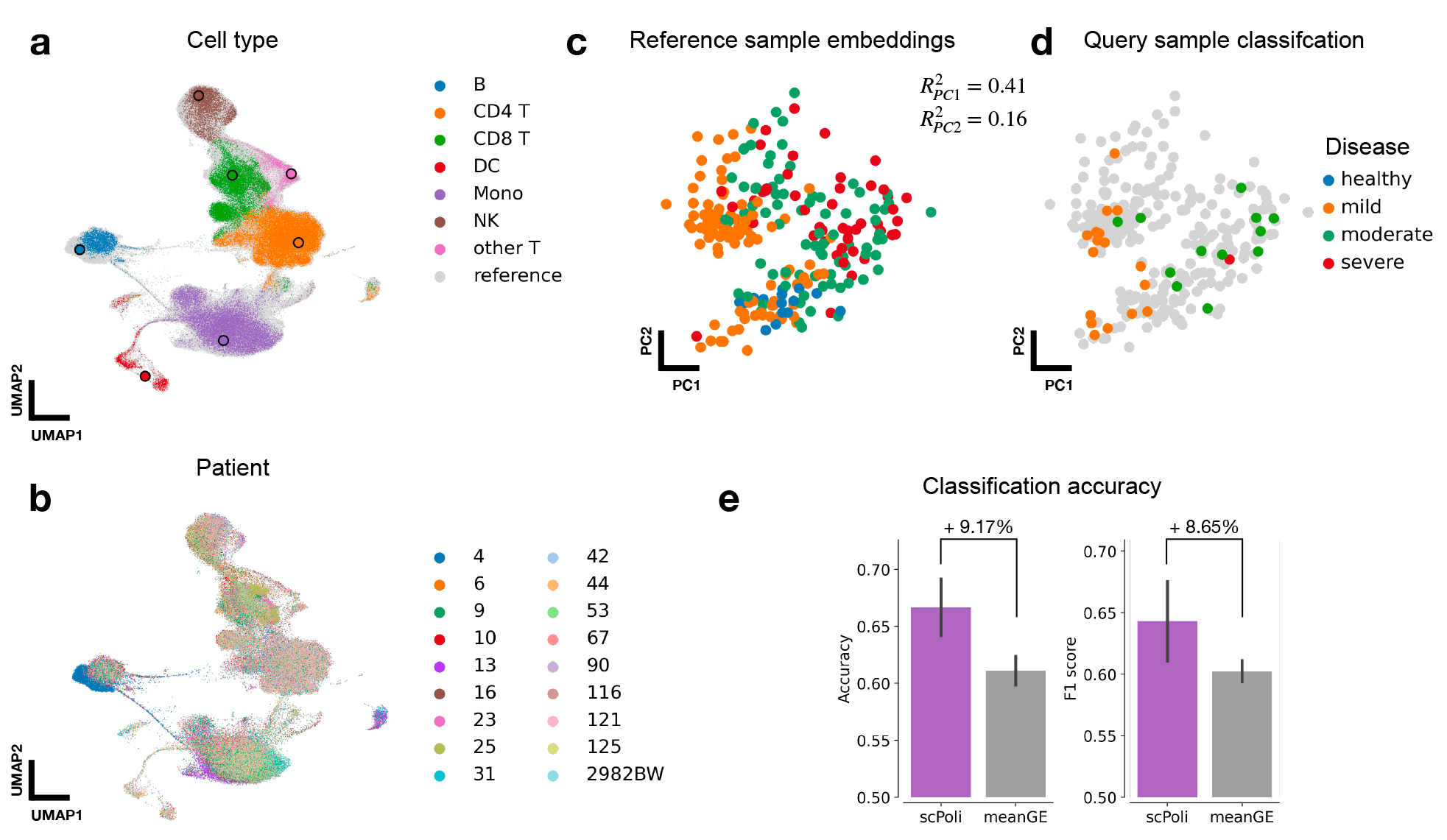
scPoli allows classification of disease state for unlabeled samples. **(a)** UMAP of Su *et al*. dataset after integration. Unlabeled cells are shown colored by the predicted cell type. Labeled cell type prototypes are shown with bigger dots with a black border. **(b)** UMAP of the integrated dataset colored by patient. Data from 30 random samples out of 270 is shown to simplify the visualization. **(c)** Principal component analysis of the labeled sample embeddings obtained with scPoli colored by disease state. **(d)** Unlabeled sample embeddings are shown in PCA space colored by their predicted disease state on top of the reference sample embeddings in grey. **(e)** Comparison of classification accuracy and F1 score obtained on scPoli embeddings and average gene expression vectors for each sample (pseudobulks).

The sample-level metadata are organized in four classes: healthy, mild, moderate and severe. The sample embeddings of the reference data showed variation associated with this phenotypic covariate in the two first principal components (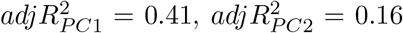 obtained with an ordinal regression) (**Fig. 4 c**). We therefore proceeded to classify the disease state of the query samples using a kNN classifier trained on the reference sample embeddings. This classification yielded an accuracy of 73%. We compared the performance and stability of this sample-level label transfer against the performance obtained with the same classifier trained on vectors of average gene expression per sample. We did so by splitting the training data in labeled and unlabeled in a 5-fold crossvalidation setting. When we compared the accuracy and F1 score obtained by a classifier trained on scPoli sample embeddings we observed that these were overall consistently better (9.17% median improvement in accuracy and, 8.65% median improvement in F1 score) than metrics obtained using training data pseudobulks (**Fig. 4 e**).

We thus believe that, in the case of a collection of controlled experiments with similar technical factors, scPoli can be used to automatically classify unlabeled data at both cell type and sample levels. This can be useful especially for disease classification as shown here, but also for other covariates of biological interest that display strong variation in the sample embedding space.

### scPoli sample embeddings support experimental design in data integration workflows

To understand the relationship between technical (e.g. assay, cohort, etc.) and phenotypic factors in scPoli’s sample embedding we integrated another COVID-19 dataset consisting of 222,003 cells (Schulte-Schrepping *et al*. (2020)) [38], with healthy and disease samples *(n* = 99) from 65 patients in two different cohorts (**Fig. 5 a, b**) obtained from a multitude of experiments with different technical properties. When we examined the sample embedding, we observed the major sources of variation to be related to technical factors rather than phenotypic ones, suggesting that these factors play a bigger role behind undesired batch effects. Indeed, the first two principal components of the sample embedding were explained by the experiment 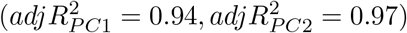 (**Fig. 5 c**) and the cohort from which the samples were obtained 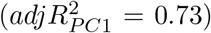 (**Fig. 5 d**), rather than the disease state 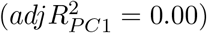 (**Fig. 5 e**). Nonetheless, we could find association with disease state, but only in further principal components of the sample embedding 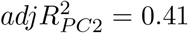, **Fig. 5e f**).

**Figure 5:**
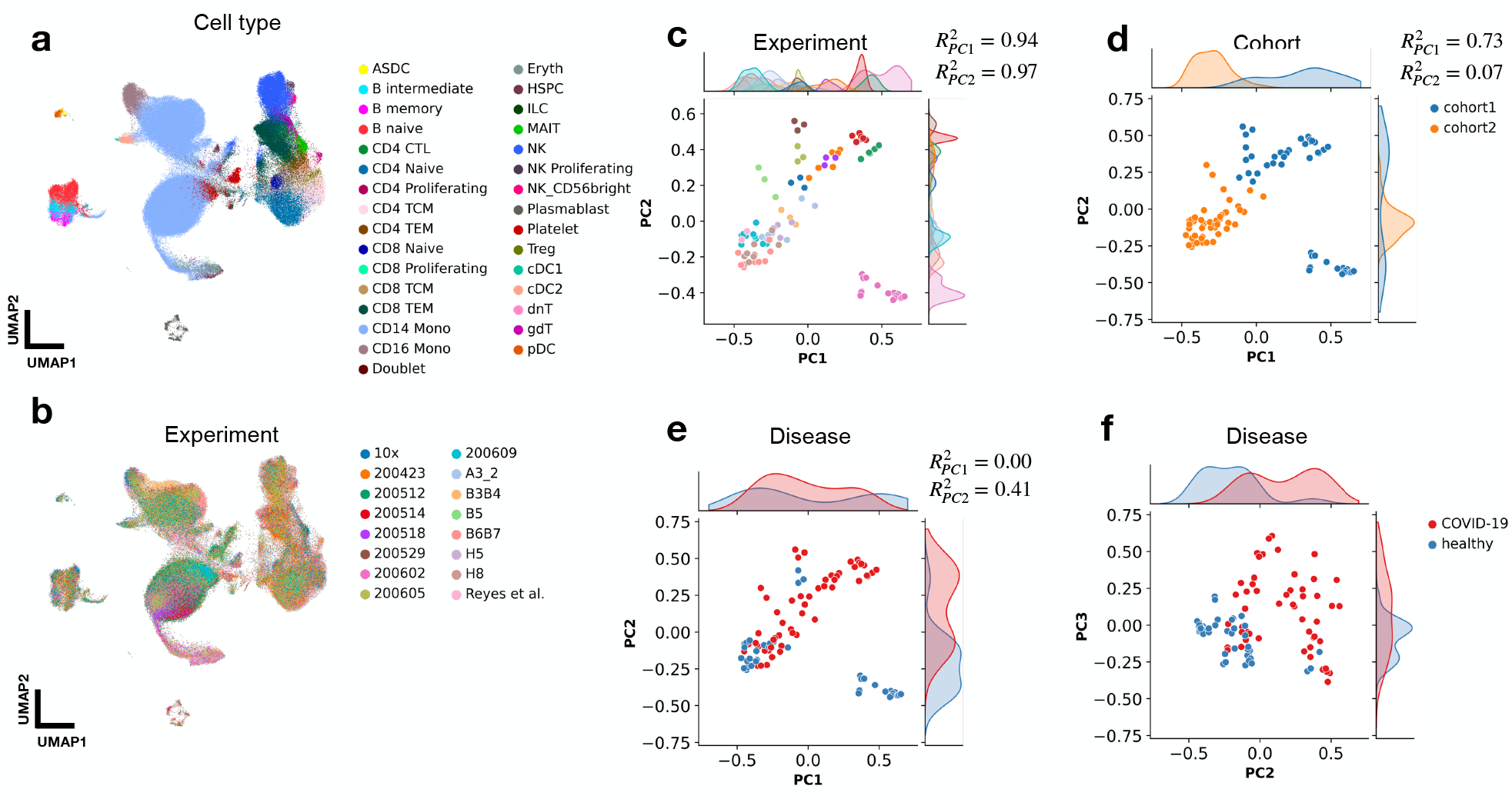
scPoli sample embeddings capture technical variation and can guide data integration workflows. **(a, b)** UMAPs of Schulte-Schrepping *et al*. dataset consisting of healthy and COVID-19 PBMC samples after sample-level integration. Cells are colored by cell type and experiment respectively. **(c)** Sample embeddings obtained with scPoli colored by experiment (legend shared with (b)), **(d)** cohort and **(e, f)** disease. For each covariate the association with the first and second principal components is displayed.

These analysis suggest that, while in more focused studies where technical factors are controlled and kept consistent across samples, biological signals represent the main source of sample-level heterogeneity; in bigger scale collections of data with a variety of technical factors, these variations will dominate. This led us to speculate that, since sample embeddings identify major sources of variations in the data, they could also guide the choice of the covariate to use as the batch to integrate in a data integration workflow. We thus proceeded to integrate the data using other two scPoli models conditioned on the covariates that showed association with the first principal component of the embedding space (experiment and cohort). We observed that the integration yielded by the cohort- and experiment-level models displayed very similar overall quality of integration despite the drastic reduction in the amount of conditions to integrate (2 for the cohort covariate and 16 for experiment, compared to 99 for the sample covariate) (**Supp. Fig. 4)**. The model trained to integrate samples yielded a mean integration score of 0.70, while those trained on the experiment and cohort covariate obtained 0.69 and 0.66 respectively. We believe this demonstrates a potentially important use-case of this novel sample-level representation. Revealing the main sources of undesired batch effects can in fact lead to leaner data integration workflows with fewer conditions, but can also potentially improve the quality of the integrated cell representations by selecting the most appropriate batch covariate.

### scPoli allows to efficiently construct a PBMC atlas with thousands of samples

We further leveraged scPoli to build a cross-disease, cross-tissue PBMC atlas comprising roughly 7.8 million cells from 2,375 samples, 1,977 subjects and 25 datasets. We obtained the integrated cell representation (**Fig. 6 a, b**, **Supp. Fig. 5 a**) and the sample embeddings which we analysed to examine the dominant sources of variations across samples. We used sample embeddings with a dimensionality of 20 and we observed that, while most of the variance was explained by the first principal component (PC) of this space, substantial signal was still present in the further PCs, suggesting that scPoli makes use of the full dimensionality of this space to encode sample-related information useful for batch correction (**Supp. Fig. 5 b**). We found the first principal component to be mainly associated with the dataset of origin of the sample (**Fig. 6 c**, **Supp. Fig. 5 c, d**). We used a linear model to quantify this association, which yielded an adjusted R^2^ of 0.97. We also observed a strong association with sequencing assay (adjusted R^2^ = 0.86) (**Fig. 6d, Supp. Fig. 5g, h** and moderate association with disease phenotype (adjusted R^2^ = 0.59) (**Fig. 6e**, **Supp. Fig. 5 e, f**). When we looked at how information such as sex and ethnicity mapped onto the sample embedding obtained with scPoli, we observed no clear patterns in the embedding space (**Supp. Fig. 5 i, j, k, l, m, n**). We compared the structure in scPoli sample embeddings with that obtained on vectors of average gene expression by sample, and observed that, while some of the patterns showed similarities, scPoli was more sensitive to differences between datasets and preserved more structure overall (**Supp. Fig. 6**).

**Figure 6:**
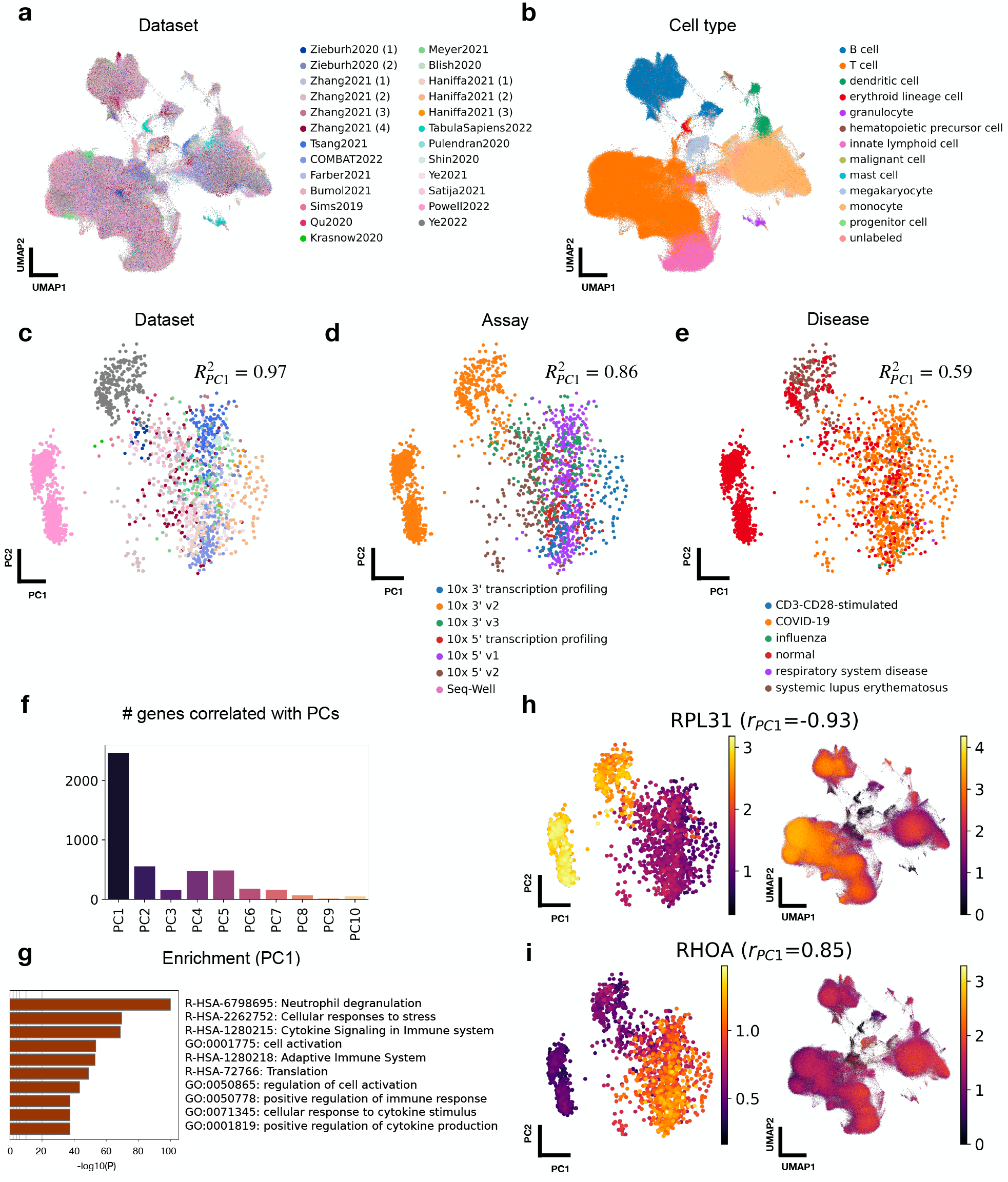
scPoli enables population-level integration of 7.8 million cells from 2,375 PBMC samples. **(a)** UMAP of the PBMC atlas after integration. Colors show different datasets of origin. In **(b)** the same UMAP is colored by cell type. **(d)** Sample embeddings projected onto the two first principal *c*omponents and colored by dataset of origin (legend shared with (a)), **(d)** assay and **(e)** disease. The displayed R^2^ is the adjusted R^2^ obtained by fitting a linear model on the first principal components and the covariates. **(f)** Number of genes significantly (*p* < 0.01 and |*r*| > 0.3) correlated with the principal components of the sample embedding space. **(g)** Biological process and pathway enrichment analysis of the genes found to be significantly correlated with PC1. **(h)** *RPL31* and **(i)** *RHOA* expression patterns in the sample embedding space (left) and cell embedding space (right). These genes were respectively amongst the most negatively and positively correlated ones with PC1.

In order to understand whether the sample embeddings reflected any gene expression patterns across samples, we computed the Pearson correlation between the mean expression of each gene in the various samples and the principal component scores of the embeddings. We thus obtained lists of significantly correlated genes with each PC (*p* < 0.01) and filtered them for coefficients of determination larger than 0.3 in absolute value. We found the biggest number of correlated genes with the first principal component (PC). Interestingly, this number did not decrease regularly going further through the PCs, and we found PC2, PC4 and PC5 to also have a substantial amount of correlated genes (**Fig. 6 f**). When we looked at which gene were most strongly correlated with PC1, we observed a strong presence of ribosomal genes: 14 out of the 15 top negatively correlated genes. This was reflected also in a general association of PC1 with the mean ribosomal gene fraction of each sample (**Supp. Fig. 7 a**). On the other hand, we did not observe a clear association with mitochondrial gene fraction (**Supp. Fig. 7 b**). We performed a biological process and pathway enrichment analyses of the genes correlated with PC1 and found terms related to the immune and stress response, cytokine signaling, neutrophile degranulation and cell activation (**Fig. 6 g**). The top negatively correlated gene with PC1 was *RPL31* (*r* = −0.93) which is a ribosomal gene involved in the cellular response to stress, the second top positively correlated gene, on the other hand, was *RHOA* (*r* = 0.85), a gene involved in the immune response and that we observed to be more highly expressed in disease samples (**Supp. Fig. 7 c**). These findings reflect the associations found with both technical and phenotypic covariates and PC1. When we looked at the patterns of expression of these genes in the sample and cell embeddings spaces we observed that scPoli successfully mixed these signals in the integrated cell representation, but also offered the unique feature of exploring them in the sample embedding space where they were preserved (**Fig. 6 h, i**). A similar enrichment analysis with PC2 and PC4-correlated genes revealed terms related RNA and DNA metabolism in the first case (**Supp. Fig. 7 d**), and response to stress and cytokine production in the second (**Supp. Fig. 7 f**). We did not perform this analysis with PC3, due to the low number of correlated genes. We show the expression patterns in both cell and sample embedding spaces of genes associated with PC2 *(SSR2*) and PC4 *(TSLPY2*) in **Supp. Fig. 7 f, g**.

We believe that the multi-scale representation obtained by scPoli could represent a useful tool for researchers to understand which genes drive batch effects the most or are affected by technical factors in the data generation process. scPoli allows the user to explore between-sample variations in gene expression and technical or phenotypic factors, while also providing the desired integrated cell representations needed for atlas building.

## Discussion

We have presented *scPoli*, a deep conditional generative model for data integration, label transfer and query-to-reference mapping. scPoli learns representations of the input data at two different scales by learning cell-level and sample-level embeddings. This enables multi-scale analyses whereby the user can explore sample information in a novel latent space, while still having access to an integrated single-cell object. By freezing the weights of the model and learning new conditional embeddings, scPoli is able to quickly map newly generated data onto a previously built reference. This step requires learning only a few free parameters, and is therefore not as computationally demanding as re-integrating from scratch.

In benchmarks we have demonstrated that scPoli is competitive with the most used and best performing methods for data integration and label transfer. Thanks to the use of cell type prototypes, scPoli consistently preserved biologically information better than other methods, while achieving comparable results in batch correction. Moreover scPoli performs label transfer in a privacy-aware fashion without need for the reference data, but only a trained reference model. We illustrated the features of our model by integrating one of the latest atlases of lung, the Human Lung Cell Atlas (HLCA). scPoli outperformed the model used in the original study in data integration and yielded a sample-level latent space which reflected similarities between different samples. We also showcased our model’s ability to be extended at a second stage by integrating an unlabeled query dataset of healthy samples and one of cancer samples. In these tasks scPoli achieved state-of-the-art label transfer performance and unknown population detection.

To understand better the information captured by the novel sample embeddings and potential use cases we investigated three further datasets. Our findings suggest that in smaller scale studies scPoli reveals phenotypical sources of variation and can enable multi-scale classification of both cell types and samples. Nonetheless as the complexity and number of samples increase the sample embeddings obtained with our model are more likely to reflect variations of technical nature. Experimental protocols and other technical differences are in fact more likely to occur as the number of samples to integrate increases. In these cases scPoli’s sample embeddings can help identify the main sources of technical variation which are driving the batch effects. This can be used to guide data integration workflows by identifying the most appropriate covariates to use as batch condition and to discover gene expression patterns across samples associated with batch effects, technical and phenotypic factors.

scPoli is integrated in the *scArches* package, which collects various models for data integration and query-to-reference mapping. This package includes also important utilities for model sharing.

scPoli, like other methods for data integration which leverage conditional variational autoencoders (CVAE), provides the user with a lower-dimensional single-cell integrated object and not a corrected count matrix. These models offer the possibility to reconstruct a representation of the data in gene space, but such approaches result in artifacts similar to denoising. This can be a limitation when gene-specific information is needed for downstream analyses with this integrated matrix. Moreover, the quality of the integration will be a function of the number of samples that can be used in the reference building step. As the model integrates samples with different technical or phenotypical characteristics it does a better job at regressing out the batch effects.

We found that the use of prototypes improves the preservation of biological information. We use euclidean distances from these prototypes and a latent cell embedding to transfer labels and yield an uncertainty associated with it. While latent representation obtained with VAEs are learned on smoother manifolds than vanilla autoencoders, a linear approximation over the whole extent of this manifold remains suboptimal. This becomes especially relevant for our uncertainty proxy, whose distribution can vary significantly in different scenarios. Therefore, we recommend users to visualize these distributions and choose the best threshold for detecting unknown cells manually. We believe the use of geodesics or other more sophisticated distances can lead to an important improvement in label transfer accuracy [39], but also in other methods which leverage distances or vector arithmetics in latent space [16].

We recommend care when interpreting the sample-level representations obtained with scPoli. The main sources of variation between samples will change across datasets. Different covariates are likely to explain these variations in different datasets. This will determine which are the most sensible use cases for sample embeddings.

We believe scPoli will be useful as a tool for data integration and query-to-reference mapping given its improvements in conservation of biological signal. Most importantly, we expect scPoli’s novel sample-level embeddings to provide researchers with a new point-of-view over large-scale datasets, and pave the way to novel multi-scale analyses which investigate and link patterns at the cell and sample level. Single cell atlassing is entering the stage of population level studies, which implies the need for models across this level of variation. With scPoli we believe we offer an efficient tool for population level data integration and mapping.

To conclude, we expect scPoli to accelerate the efficient construction and annotation of large-scale reference atlases, and help model reuse for the broad set of atlas users.

## Code availability

The software is available as part of https://scarches.readthedocs.io. The code to reproduce the results is available at http://github.com/theislab/scPoli_reproduce.

## Data availability

All datasets analyzed in this manuscript are public and have been published in other papers. We have referenced them in the manuscript and when necessary made them available at http://github.com/theislab/scPoli_reproduce.

## Author Contributions

M.L. conceived the project with F.J.T.. M.L, C.D. and M.W. designed the algorithm. C.D. and M.W. implemented the algorithm with contributions from A.A.M. C.D., M.W., L.Z. curated the datasets used for the analyses in the paper. C.D. and S.H.Z. performed experiments and analyses. C.D. ran benchmarking experiments. All authors contributed to the manuscript. M.L. and F.J.T. supervised the project.

## Acknowledgments

We thank Lisa Sikkema for the valuable feedback on our work with the HLCA dataset. We thank Sergey Rybakov for helping integrate our software to the scArches package. M.L. acknowledges financial support from the Joachim Herz Stiftung. FJT acknowledges support by the the Helmholtz Association’s Initiative and Networking Fund through Helmholtz AI [ZT-I-PF-5-01].

## Competing interests

F.J.T. consults for Immunai Inc., Singularity Bio B.V., CytoReason Ltd, and Omniscope Ltd, and has ownership interest in Dermagnostix GmbH and Cellarity. C.D. is a part-time employee of Immunai, Inc.

## Methods

### scPoli

scPoli builds upon Conditional Variational Autoencoders (CVAEs[15, 40]), but with an important modification. While in a standard CVAE different conditions *s*_1_,…, *s_N_* are represented by means of a fixed one-hot-encoded vector which is concatenated to the input *x*, we use learnable embeddings 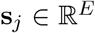 of fixed dimensionality *E* to represent each category we want to regress out. The learning objective for this network is akin to that of a standard CVAE, but learning the embeddings *S* during training:

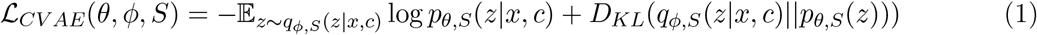

### Prototypes for label transfer

After a pre-training step we initialize *k* labeled prototypes, where *k* is the number of distinct cell types present in the reference datasets. We compute these prototypes 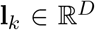 where *D* is the dimensionality of the latent space of the CVAE model, by computing the average latent representation of the data points belonging to each particular cell type:

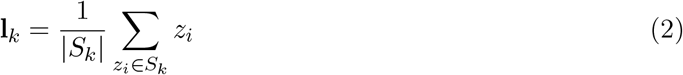

Where *z_i_* is the latent representation of cell *i, k* is the cell type label and *S_k_* is the set of cells belonging to cell type *k*.

### Prototype loss

After pre-training, we add another term to the training objective which we call prototype loss. This term has the objective of pulling together samples belonging to the same cell type towards their correspondent prototype in latent space.

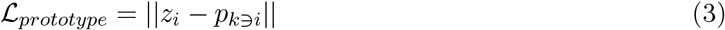

Where *p_k∋i_* denotes the prototype of the cell type class to which *z_i_* belongs.

### Reference mapping

We map new studies onto a reference model by freezing the weights of the encoder and decoder net-works as prevously proposed by scArches[21] and adding a set of M randomly initialized learnable embeddings implemented using ‘torch.nn.Embedding’ in PyTorch, which are the only learnable parameters at this stage. The learning objectives are the same as those used during reference building.

### Cell type label transfer and uncertainty quantification

After the model is trained on query data, we assign to query cell i with latent representation *z_i_* the label *K_i_* of the closest prototype p in latent space:

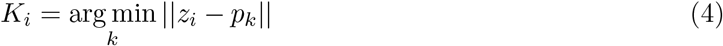

The minimum distance between to reference prototype is used as a proxy for uncertainty for unknown cell type detection.

### Training

The training steps for a standard workflow of scPoli are the following:

- Reference building

– Pre-training: initialize N embeddings, where N is the number of studies present in the reference, optimize 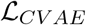 on the reference dataset;
– Fine-tuning: initialize cell type prototypes and optimize 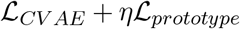, where *η* is a hyperparameter, store learned prototypes with the model;
- Reference mapping

– Pre-training: freeze the weights of the encoder and decoder networks from the reference model, initialize M additional learnable embeddings where M is the number of batches in the query, optimize 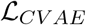 on the query data;
– Fine-tuning: initialize unlabeled prototypes in the query dataset after unsupervised clustering of the latent representation of the data, optimize 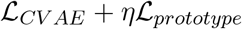, if all cells in the query are unlabeled this learning objective is reduced to 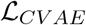.

### Hyper-parameters and training

For each dataset, we performed a hyperparameter search over the most important parameters of our model. These include the depth of the encoder and decoder architecture, as well as the weight *η* for the prototype loss. We fixed as many hyperparameters as possible to keep the computational overhead withing a reasonable limit. We also found that the model performs consistently across hyperparameter choices. We selected the hyperparameters that yielded the best integration scores averaged across the datasets used for benchmarking. Thus, we use **identical architecture and hyperparameters** for all benchmarks shown in the paper.

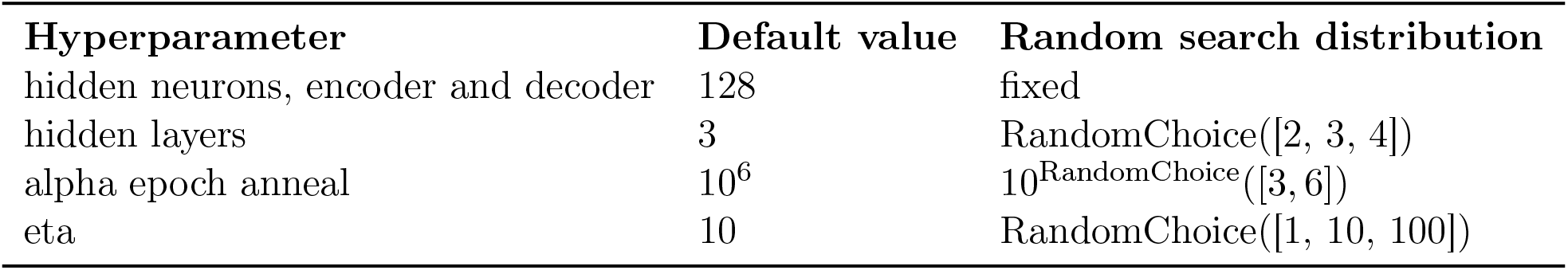

### Benchmarks

#### Integration methods

We benchmarked the data integration performance of our model against various state-of-the-art methods. These include:

- **scArches scVI (v0.5.3)**: we ran the model using default parameters;
- **scArches scANVI (v0.5.3)**: we ran the model using default parameters;
- **Seurat v3 (v4.0.3)**: we followed the tutorial (https://satijalab.org/seurat/archive/v3.0/integration.html) and used supervised PCA for reducing the data dimensionality to 50;
- **Symphony (v0.1.0**: we followed the vignette (https://cran.r-project.org/web/packages/symphony/vignettes/quickstart_tutorial.html) and ran the model using default parameters;

#### Cell type classification methods

We benchmarked the performance on label transfer and cell type classification against the following methods:

- **scArches scANVI (v0.5.3)**: we ran the model using default parameters;
- **MARS**: we ran the model using default parameters;
- **Seurat v3 (v4.0.3)**: we followed the vignette (https://satijalab.org/seurat/archive/v3.0/integration.html) and ran the model using default parameters;
- **SVM**: we fit a *LinearSVC* object from *scikit-learn* (v0.24.2) on the reference data;

### Metrics

We quantified the quality of the data integration using the following metrics from the *scIB* (v 1.0.0) package and Luecken *et al*. We provide a short explanation of each metric, for more details we suggest to read [6].

#### NMI

NMI (Normalized Mutual Information) measures the overlap between two different clusterings. This score is used between the cell type labels and the clusters obtained via unsupervised clustering of the dataset after integration. The score is bounded between 0 and 1, with 1 denoting a perfect match between the two clusterings and 0 the absence of any overlap.

#### ARI

ARI stands for Adjusted Rand Index. The raw Rand Index scores the similarity between two clusterings, it considers both correct overlaps and correct disagreements in the computation. This score is computed between the cell type labels and the clusters obtained on the integrated dataset. The score spans from 0 to 1, with 1 representing a perfect score.

#### Cell type ASW

ASW (Average Silhouette Width) is a measure of the relationship between the within-cluster distances of a cell and the between-cluster distances of the same cell to the closest cluster. This metric can range between −1 and 1. Values of −1 indicate total misclassification, values of 0 imply overlap between clusters and values of 1 occur when clusters are well separated. We use two versions of this scores, one computed on cell-type labels, and a modified version to quantify batch mixing (see ASW batch below). The score is scaled to have values between 0 and 1 using the following equation:

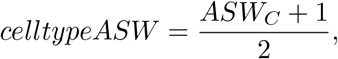

where C represents the set of all cell type labels.

#### Batch ASW

The batch ASW quantifies the quality of batch mixing in the integrated object. We obtain it by computing the ASW but on batch labels instead of cell type labels. Scores of 0 are indicative of good batch mixing, while any deviation from this score is the result of batch effects. In order to have a metric bound between 0 and 1, the following transformation is applied:

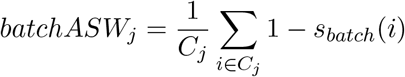

Here *C_j_* is the set of cells with label *j* and |*C_j_*| is the support of the set. The final score is obtained by averaging the batch ASW values obtained for each label:

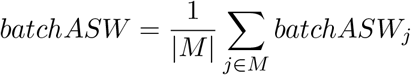

with M being the set of unique cell labels.

#### Isolated label F1

Isolated labels are defined as cell type labels which are found in the smallest number of batches. We aim to determine how well these cell types are separated from the rest in the integrated data. To do so, we find the cluster containing the highest amount of cells from such isolated labels, and we then compute the *F*_1_ score of the isolated label against all other labels within the cluster. We use the standard *F*_1_ formulation:

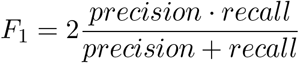

Once again, scores of 1 represent the desirable outcome in which all cells from the isolated labels are grouped in one cluster.

#### Isolated label silhouette

This is a cell type ASW score but computed only on the isolated labels.

#### Principal component regression (PCR)

This metric uses principal component regression to quantify the variance contribution of the batch covariate per principal component. This is done by multiplying the variance explained by the *i*-th PC with the *R*^2^ between the covariate of interest and the same PC. This is then summed across all PCs:

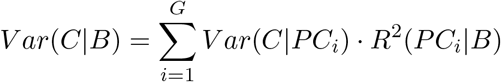

#### Graph connectivity

This metric measures whether a kNN graph (G) computed on the integrated object connects cells that fall within the same cell type. It is bound between 0 and 1, with 0 indicating a graph where all cells are unconnected and, and 1 occurring when all cells of the same cell type are connected in the integrated output. For each cell type label a subset graph *G*(*N_c_*, *E_c_*) is computed, the final score is obtained using the following formulation:

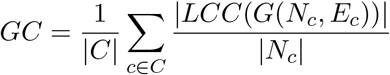

where *C* is the set of unique cell type labels, |*LCC*()| is the number of nodes in the largest connected component in the kNN graph and |N_c_| is the number of nodes in the graph.

### Datasets

#### The Human Lung Cell Atlas (HLCA)

We obtained the HLCA core dataset from the authors of the study [4]. The dataset consists of 584,884 lung cells from 166 samples and 107 subjects. Gene expression is subset to 2,000 selected genes which we used for model training. Different levels of cell type annotation and sample and patient metadata are curated and available.

#### PBMC Atlas

The atlas contains 7,800,850 PBMC cells from 2,375 samples, representing cells from 25 datasets, 1,977 healthy or diseased donors. For model training and analysis 10,000 HVGs were selected. A coarse annotation consisting of 14 cell types was used for initializing scPoli’s prototypes. The data and metadata were curated by scientists at the Chan-Zuckerberg Initiative and collected from [38, 41–58].

#### Schulte-Schrepping et al. dataset

This is a published [38] PBMC dataset of 65 COVID-19 patients and healthy controls. The dataset contains 99 samples and 222,003 cells, and was downloaded as part of the Fredhutch COVID-19 collection available at https://atlas.fredhutch.org/fredhutch/covid/. For model training and analysis 4,000 HVGs were used.

#### Su *et al*. dataset

This is a published [37] PBMC dataset of 129 COVID-19 patients and 16 healthy controls. The dataset contains 270 samples and 559,517 cells, and was downloaded as part of the Fredhutch COVID-19 collection available at https://atlas.fredhutch.org/fredhutch/covid/. For model training and analysis 2,000 HVGs were used.

### Benchmark datasets

All datasets used for benchmarking were obtained from https://github.com/theislab/scArches-reproducibility unless specified otherwise. Count data were used for all datasets.

#### PBMC

The PBMC dataset used for benchmarking was obtained from (https://doi.org/10.6084/m9.figshare.12420968)[6]. The dataset contains 32,484 cells from 4 studies and 16 cell types. The data was subset to the 4,000 most highly variable genes before further analysis.

#### Pancreas

The data contains 16,382 pancreas cells from 8 different batches. The cells are annotated and assigned to 14 cell types. 4,000 HVGs were used for model training and analysis.

#### Brain

The mouse brain dataset consists of 332,129 cells and 4 batches. 10 cell types are present. The data were subset to the 4,000 most highly variable genes before further analysis.

#### Scvelo

The data consists of 25,919 cells in 4 batches and 14 cell types. 4,000 highly variable genes were selected for downstream analyses.

#### Tumor

The tumor atlas was obtained from [59] (https://zenodo.org/record/4263972). The dataset is a collection of 14 studies on various types of cancer. It contains 317,111 cells annotated in 25 cell types. We selected 4,000 HVGs for model training.

#### Lung

The lung data were obtained from [6] (https://doi.org/10.6084/m9.figshare.12420968). These data consist of 32,472 lung cells from 3 batches and 17 cell types. 4,000 HVGs were used for model training.

---

**Supplementary Figure 1:**
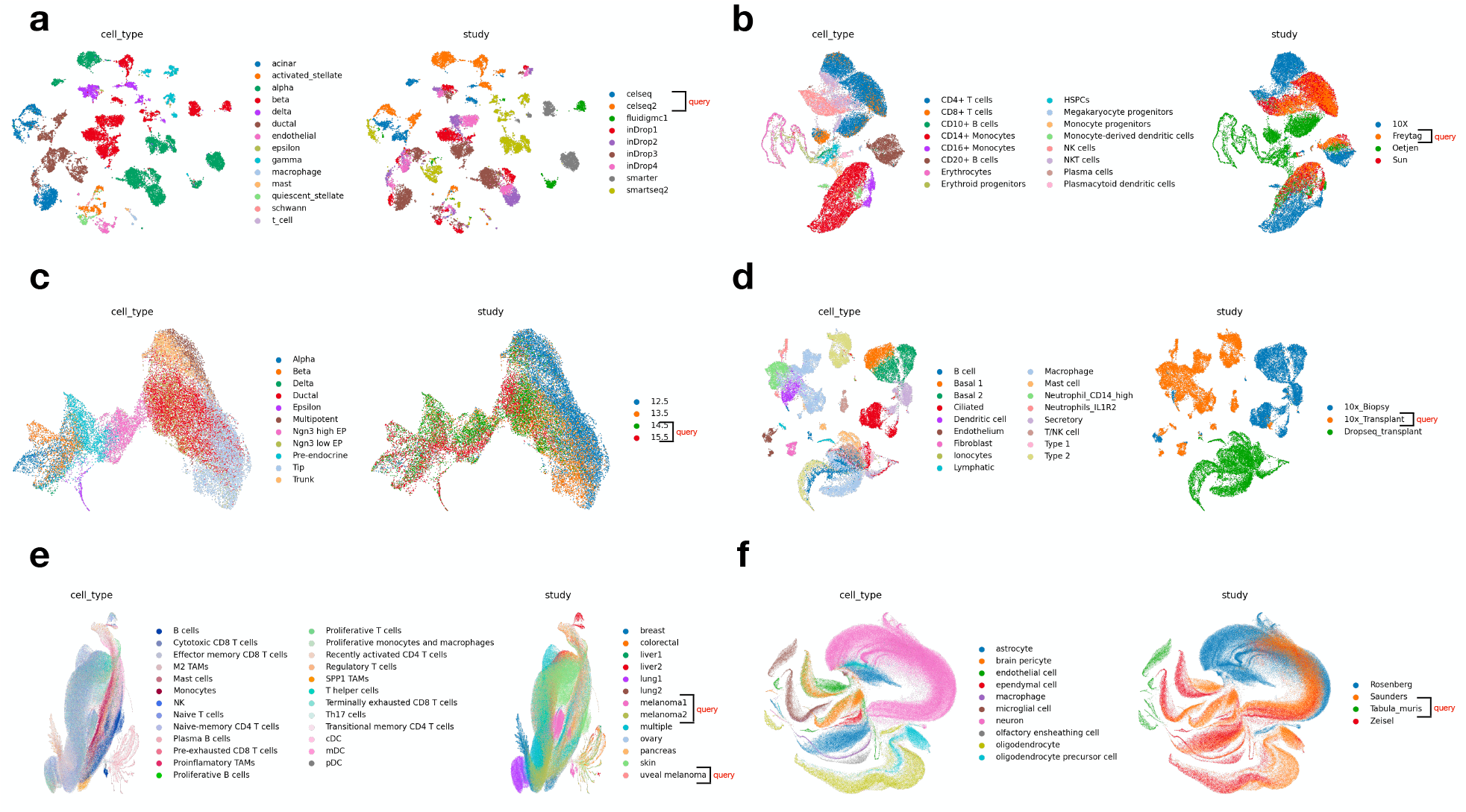
Datasets used for benchmarking. **(a)** Pancreas dataset. **(b)** PBMC dataset. **(c)** Immune dataset. **(d)** Lung dataset. **(e)** Tumor dataset. **(f)** Brain dataset.

**Supplementary Figure 2:**
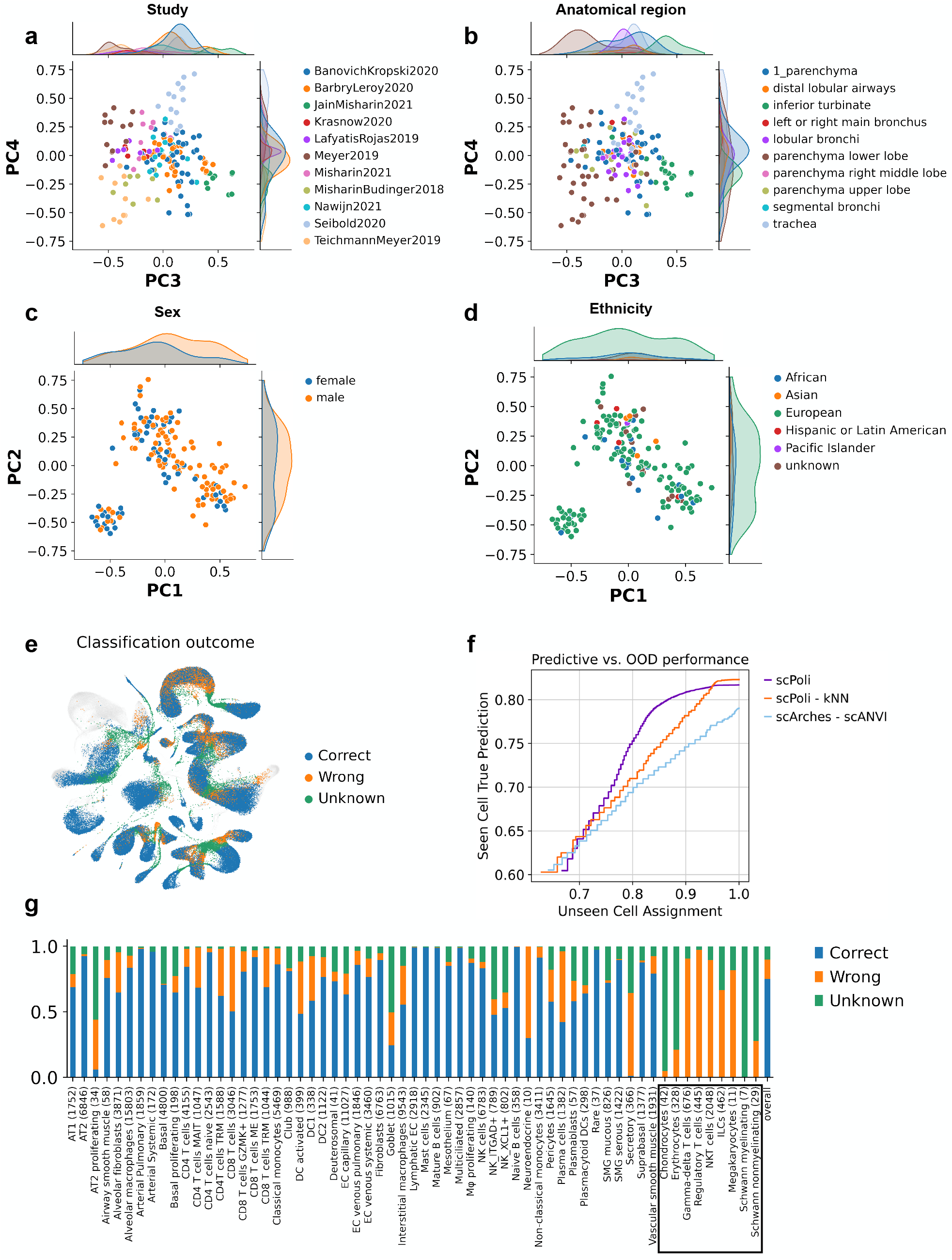
Integration and reference mapping with the Human Lung Cell Atlas. Sample embeddings colored by study **(a)**, anatomical region **(b)**, sex **(c)** and ethnicity **(d)**. **(e)** UMAP of the joint cell embedding of reference and query with classification outcome in color. Reference cells are shown in gray. **(f)** Label transfer performance comparison between scPoli, scPoli with a kNN classifier and scANVI. The accuracy is tracked across different uncertainty threshold used for unknown cell type identification. **(g)** Stacked barplot showing the accuracy of cell type classification by cell type. Cell types that were not present in the reference are marked with a box.

**Supplementary Figure 3:**
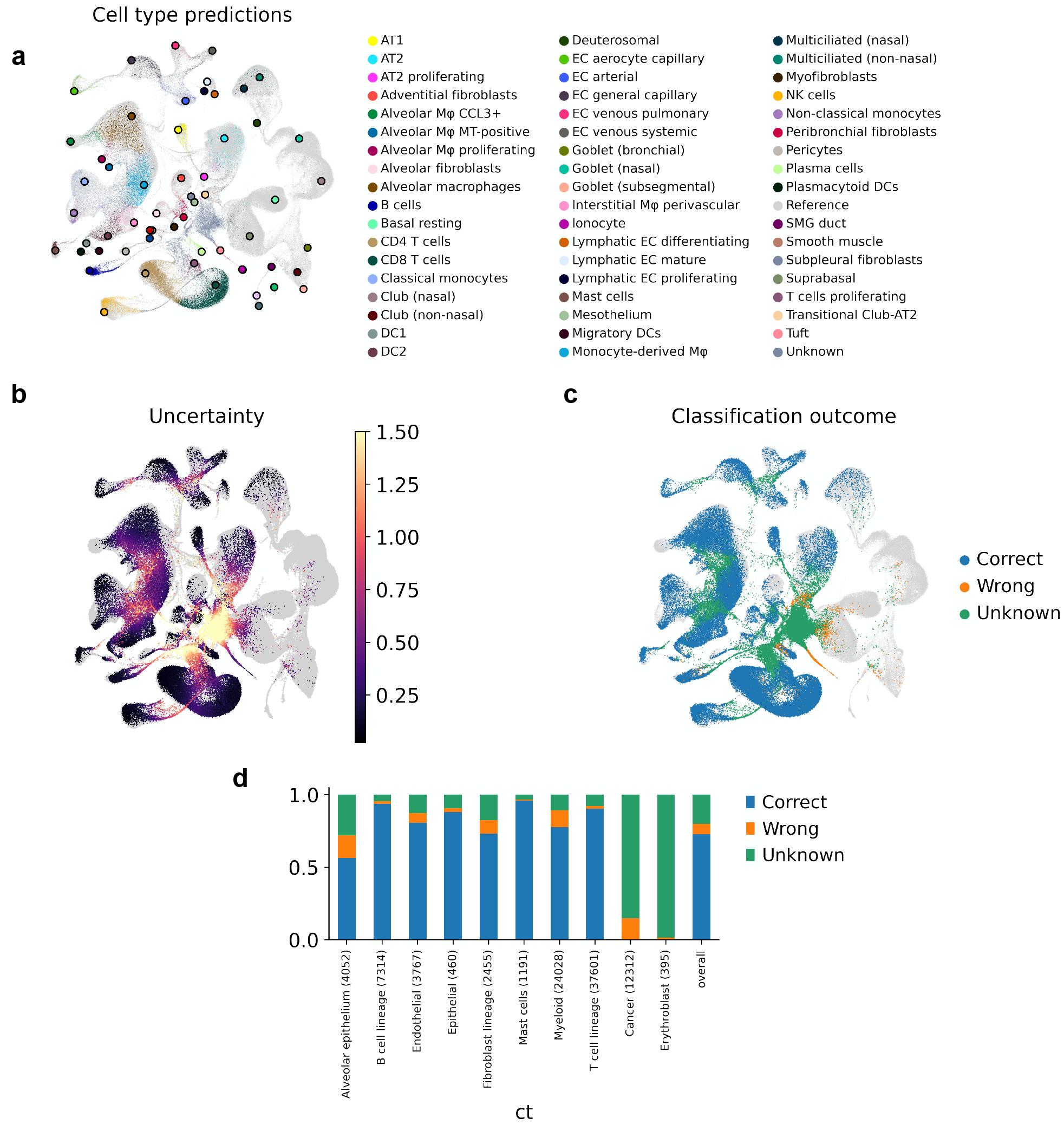
Query-to-reference mapping of a cancer dataset onto the Human Lung Cell Atlas. **(a)** UMAP of the joint query and reference datasets after query-to-reference mapping for a cancer query. Reference cells are greyed out, while query cells are colored by the predicted cell type. Reference prototype are shown as bigger dots with a black border, and are color coded by cell type. **(b)** UMAP of the integrated object with uncertainties in color. Reference cells are shown in gray. **(c)** UMAP showing the outcome of the label transfer. Reference cells are in gray. **(d)** Stacked barplot showing the accuracy of cell type classification by cell type. Cancer cells and erythrocytes were not present in the reference data.

**Supplementary Figure 4:**
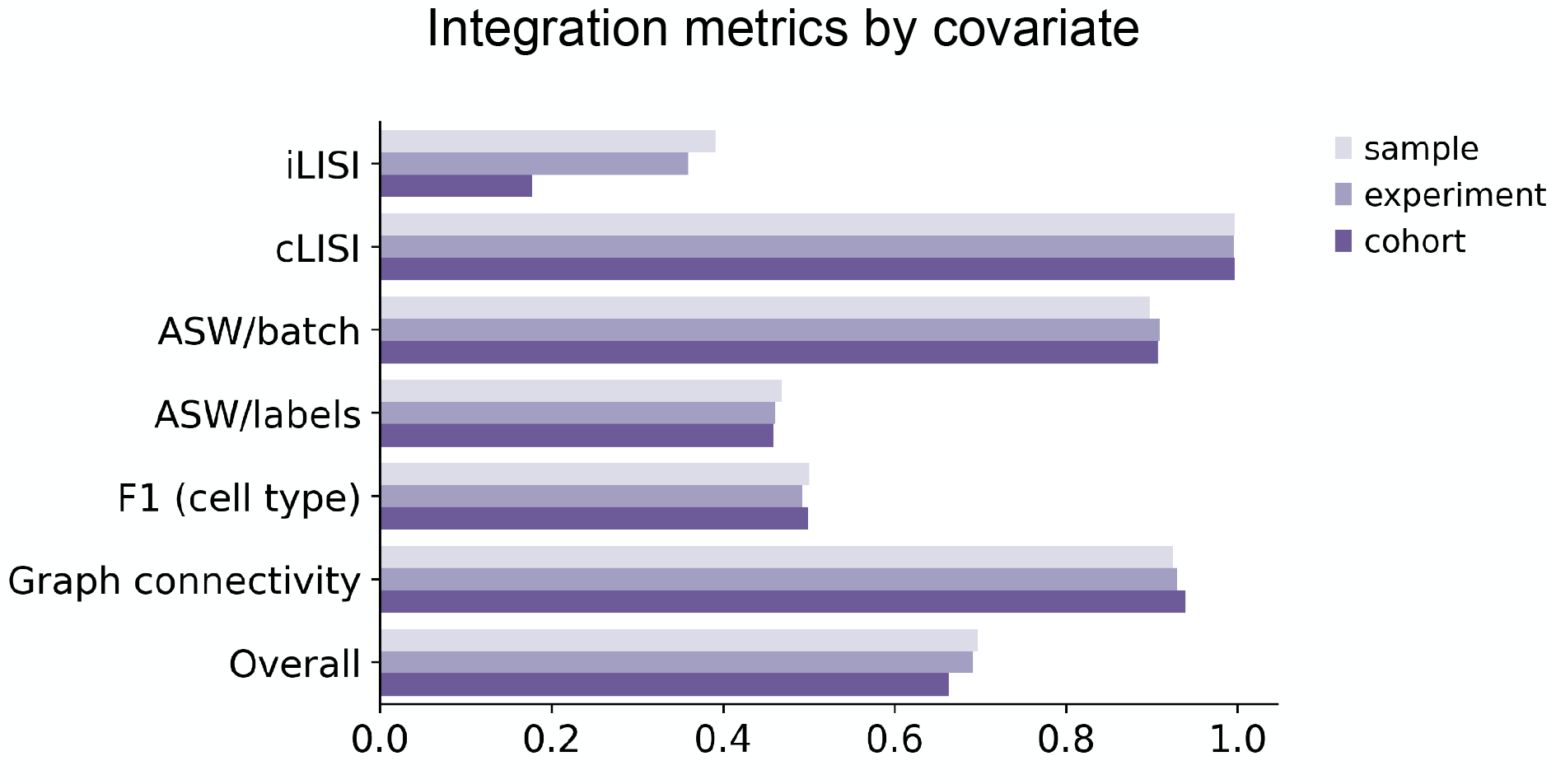
Integration metrics by covariate chosen for integration in Schulte-Schrepping et al. dataset.

**Supplementary Figure 5:**
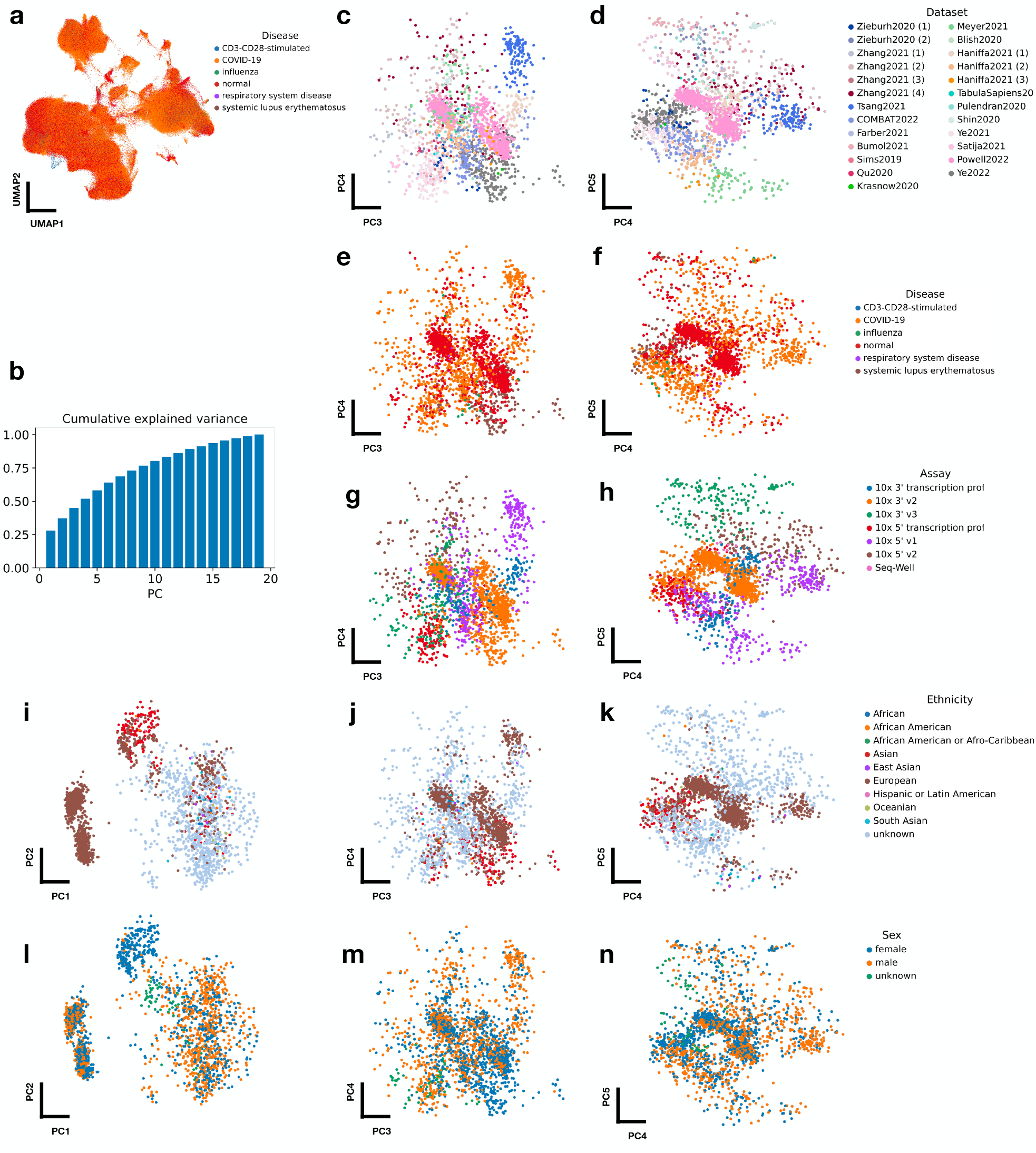
Exploration of the sample embedding space of a large-scale PBMC atlas. **(a)** UMAP of the integrated atlas, cells are colored. by disease. **(b)** Cumulative explained variance by principal component in the sample embedding space. **(c)** Samples colored by dataset of origin in PC3 and PC4, and **(d)** PC4 and PC5. **(e)** Samples colored by disease in PC3 and PC4, and **(f)** PC4 and PC5. **(g)** Samples colored by assay in PC3 and PC4, and **(h)** PC4 and PC5. **(i, j, k, l)** Samples colored by ethnicity metadata and sex metadata **(l, m, n)** in the first 5 principal components.

**Supplementary Figure 6:**
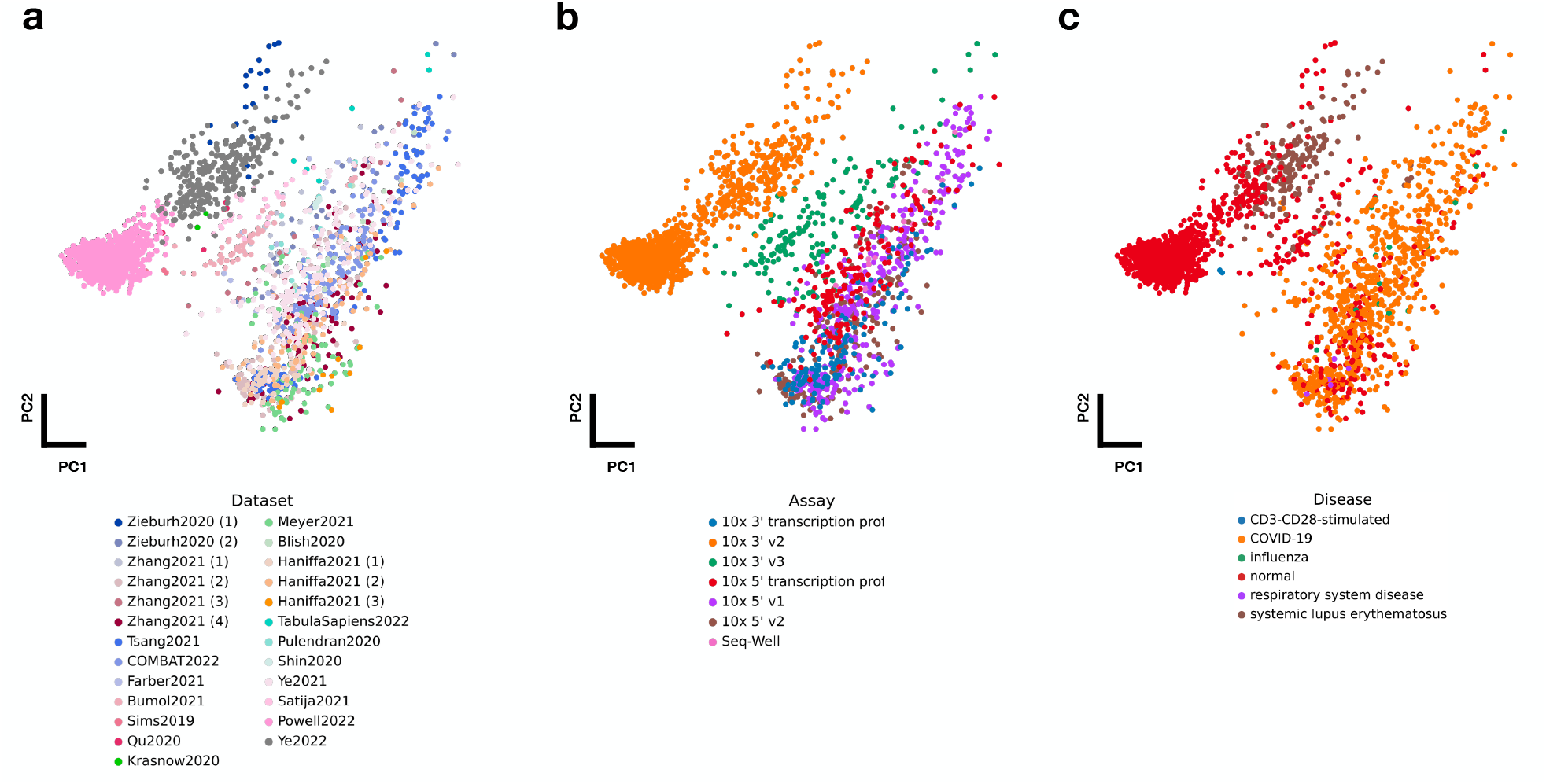
Principal component analysis of average gene expressions by sample in a large-scale PBMC atlas. First two principal components computed on the average gene expression by sample colored by **(a)** dataset, **(b)** assay and **(c)** disease state.

**Supplementary Figure 7:**
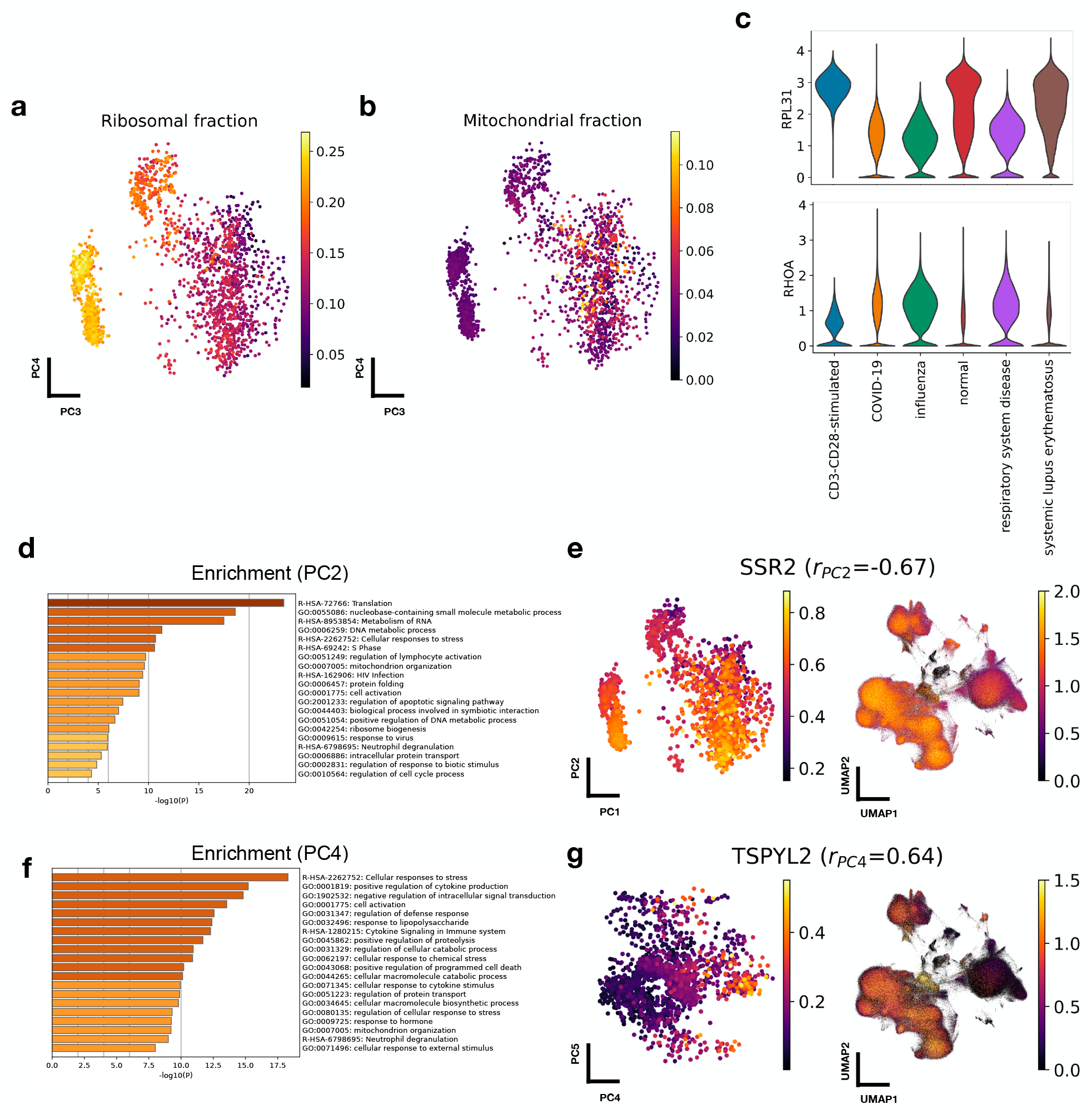
Technical factors and gene patterns across samples in a large-scale PBMC atlas. **(a)** Mean ribosomal and **(b)** mitochondrial gene fraction in the sample embedding space. **(c)** Violin plot of expression of *RPL31* and *RHOA* by disease condition. **(d)** Biological process and pathway enrichment analysis of genes significantly correlated with PC2. **(e)** *SSR2* expression patterns in the sample embedding space (left) and cell embedding space (right). **(f)** Biological process and pathway enrichment analysis of genes significantly correlated with PC4. **(g)** *TSPYL2* expression patterns in the sample embedding space (left) and cell embedding space (right).

## References

[1] Regev, A. et al. Science forum: the human cell atlas. elife 6, e27041 (2017).

[2] lcai@ caltech. edu 21 b Shendure Jay 9 Trapnell Cole 9 Lin Shin shinlin@ uw. edu 2 e Jackson Dana 9, C.-U. T. C. L. et al. The human body at cellular resolution: the nih human biomolecular atlas program. Nature 574, 187–192 (2019).

[3] Muus, C. et al. Single-cell meta-analysis of sars-cov-2 entry genes across tissues and demographics. Nature medicine 27, 546–559 (2021).

[4] Sikkema, L. et al. An integrated cell atlas of the human lung in health and disease. bioRxiv (2022).

[5] Gayoso, A. et al. A python library for probabilistic analysis of single-cell omics data. Nature Biotechnology 40, 163–166 (2022).

[6] Luecken, M. D. et al. Benchmarking atlas-level data integration in single-cell genomics. Nature methods 19, 41–50 (2022).

[7] Argelaguet, R., Cuomo, A. S., Stegle, O. & Marioni, J. C. Computational principles and challenges in single-cell data integration. Nature biotechnology 39, 1202–1215 (2021).

[8] Welch, J. D. et al. Single-cell multi-omic integration compares and contrasts features of brain cell identity. Cell 177, 1873–1887 (2019).

[9] Ritchie, M. E. et al. limma powers differential expression analyses for rna-sequencing and microarray studies. Nucleic acids research 43, e47–e47 (2015).

[10] Korsunsky, I. et al. Fast, sensitive and accurate integration of single-cell data with harmony. Nature methods 16, 1289–1296 (2019).

[11] Kiselev, V. Y., Yiu, A. & Hemberg, M. scmap: projection of single-cell rna-seq data across data sets. Nature methods 15, 359–362 (2018).

[12] Stuart, T. et al. Comprehensive integration of single-cell data. Cell 177, 1888–1902 (2019).

[13] Polański, K. et al. Bbknn: fast batch alignment of single cell transcriptomes. Bioinformatics 36, 964–965 (2020).

[14] Haghverdi, L., Lun, A. T., Morgan, M. D. & Marioni, J. C. Batch effects in single-cell rna-sequencing data are corrected by matching mutual nearest neighbors. Nature biotechnology 36, 421–427 (2018).

[15] Lopez, R., Regier, J., Cole, M. B., Jordan, M. I. & Yosef, N. Deep generative modeling for single-cell transcriptomics. Nature methods 15, 1053–1058 (2018).

[16] Lotfollahi, M., Wolf, F. A. & Theis, F. J. scgen predicts single-cell perturbation responses. Nature methods 16, 715–721 (2019).

[17] Amodio, M. et al. Exploring single-cell data with deep multitasking neural networks. Nature methods 16, 1139–1145 (2019).

[18] Lähnemann, D. et al. Eleven grand challenges in single-cell data science. Genome biology 21, 1–35 (2020).

[19] Hao, Y. et al. Integrated analysis of multimodal single-cell data. Cell 184, 3573–3587 (2021).

[20] Kang, J. B. et al. Efficient and precise single-cell reference atlas mapping with symphony. Nature communications 12, 1–21 (2021).

[21] Lotfollahi, M. et al. Query to reference single-cell integration with transfer learning. bioRxiv (2020).

[22] Michielsen, L. et al. Single-cell reference mapping to construct and extend cell type hierarchies. bioRxiv (2022).

[23] Osorio, D., McGrail, D. J., Sahni, N. & Yi, S. S. Drug combination prioritization for cancer treatment using single-cell rna-seq based transfer learning. bioRxiv (2022).

[24] Xu, C. et al. Probabilistic harmonization and annotation of single-cell transcriptomics data with deep generative models. Molecular systems biology 17, e9620 (2021).

[25] Fetaya, E., Jacobsen, J.-H., Grathwohl, W. & Zemel, R. Understanding the limitations of conditional generative models. arXiv preprint arXiv:1906.01171 (2019).

[26] Brbić, M. et al. Mars: discovering novel cell types across heterogeneous single-cell experiments. Nature methods 17, 1200–1206 (2020).

[27] Sohn, K., Lee, H. & Yan, X. Learning structured output representation using deep conditional generative models. Advances in neural information processing systems 28 (2015).

[28] Snell, J., Swersky, K. & Zemel, R. Prototypical networks for few-shot learning. Advances in neural information processing systems 30 (2017).

[29] Lotfollahi, M., Naghipourfar, M., Theis, F. J. & Wolf, F. A. Conditional out-of-distribution generation for unpaired data using transfer vae. Bioinformatics 36, i610–i617 (2020).

[30] Snell, J., Swersky, K. & Zemel, R. Prototypical networks for few-shot learning. Advances in neural information processing systems 30 (2017).

[31] Hospedales, T., Antoniou, A., Micaelli, P. & Storkey, A. Meta-learning in neural networks: a survey. arxiv preprint arxiv: 200405439 (2020).

[32] Lopez, R., Regier, J., Cole, M. B., Jordan, M. I. & Yosef, N. Deep generative modeling for single-cell transcriptomics. Nature methods 15, 1053–1058 (2018).

[33] Xu, C. et al. Probabilistic harmonization and annotation of single-cell transcriptomics data with deep generative models. Molecular systems biology 17, e9620 (2021).

[34] Stuart, T. et al. Comprehensive integration of single-cell data. Cell 177, 1888–1902 (2019).

[35] Kang, J. B. et al. Efficient and precise single-cell reference atlas mapping with symphony. Nature communications 12, 1–21 (2021).

[36] Grabski, I. N., Street, K. & Irizarry, R. A. Significance analysis for clustering with single-cell rna-sequencing data. bioRxiv (2022).

[37] Su, Y. et al. Multiomic immunophenotyping of covid-19 patients reveals early infection trajectories. BioRxiv (2020).

[38] Schulte-Schrepping, J. et al. Severe covid-19 is marked by a dysregulated myeloid cell compartment. Cell 182, 1419–1440 (2020).

[39] Engelmann, J. et al. Uncertainty quantification for atlas-level cell type transfer. arXiv preprint arXiv:2211.03793 (2022).

[40] Sohn, K., Lee, H. & Yan, X. Learning structured output representation using deep conditional generative models. Advances in neural information processing systems 28 (2015).

[41] Lee, J. S. et al. Immunophenotyping of covid-19 and influenza highlights the role of type i interferons in development of severe covid-19. Science immunology 5, eabd1554 (2020).

[42] Stephenson, E. et al. Single-cell multi-omics analysis of the immune response in covid-19. Nature medicine 27, 904–916 (2021).

[43] Szabo, P. A. et al. Longitudinal profiling of respiratory and systemic immune responses reveals myeloid cell-driven lung inflammation in severe covid-19. Immunity 54, 797–814 (2021).

[44] Yoshida, M. et al. Local and systemic responses to sars-cov-2 infection in children and adults. Nature 602, 321–327 (2022).

[45] Savage, A. K. et al. Multimodal analysis for human ex vivo studies shows extensive molecular changes from delays in blood processing. Iscience 24, 102404 (2021).

[46] Yazar, S. et al. Single-cell eqtl mapping identifies cell type–specific genetic control of autoimmune disease. Science 376, eabf3041 (2022).

[47] Guo, C. et al. Single-cell analysis of two severe covid-19 patients reveals a monocyte-associated and tocilizumab-responding cytokine storm. Nature communications 11, 1–11 (2020).

[48] Arunachalam, P. S. et al. Systems biological assessment of immunity to mild versus severe covid-19 infection in humans. Science 369, 1210–1220 (2020).

[49] Ahern, D. J. et al. A blood atlas of covid-19 defines hallmarks of disease severity and specificity. MedRxiv (2021).

[50] Travaglini, K. J. et al. A molecular cell atlas of the human lung from single-cell rna sequencing. Nature 587, 619–625 (2020).

[51] Liu, C. et al. Time-resolved systems immunology reveals a late juncture linked to fatal covid-19. Cell 184, 1836–1857 (2021).

[52] Wilk, A. J. et al. A single-cell atlas of the peripheral immune response in patients with severe covid-19. Nature medicine 26, 1070–1076 (2020).

[53] Ren, X. et al. Covid-19 immune features revealed by a large-scale single-cell transcriptome atlas. Cell 184, 1895–1913 (2021).

[54] Consortium*, T. S. et al. The tabula sapiens: A multiple-organ, single-cell transcriptomic atlas of humans. Science 376, eabl4896 (2022).

[55] Szabo, P. A. et al. Single-cell transcriptomics of human t cells reveals tissue and activation signatures in health and disease. Nature communications 10, 1–16 (2019).

[56] van der Wijst, M. G. et al. Type i interferon autoantibodies are associated with systemic immune alterations in patients with covid-19. Science translational medicine 13, eabh2624 (2021).

[57] Perez, R. K. et al. Single-cell rna-seq reveals cell type–specific molecular and genetic associations to lupus. Science 376, eabf1970 (2022).

[58] Hao, Y. et al. Integrated analysis of multimodal single-cell data. Cell 184, 3573–3587 (2021).

[59] Nieto, P. et al. A single-cell tumor immune atlas for precision oncology. Genome research 31, 1913–1926 (2021).

